# Coordinated immune dysregulation in Juvenile Dermatomyositis revealed by single-cell genomics

**DOI:** 10.1101/2023.11.07.566033

**Authors:** Gabrielle Rabadam, Camilla Wibrand, Emily Flynn, George C. Hartoularos, Yang Sun, Chun Jimmie Ye, Susan Kim, Zev Gartner, Marina Sirota, Jessica Neely

## Abstract

Juvenile Dermatomyositis (JDM) is one of several childhood-onset autoimmune disorders characterized by a type I interferon response and autoantibodies. Treatment options are limited due to incomplete understanding of how the disease emerges from dysregulated cell states across the immune system. We therefore investigated the blood of JDM patients at different stages of disease activity using single-cell transcriptomics paired with surface protein expression. By immunophenotyping peripheral blood mononuclear cells, we observed skewing of the B cell compartment towards an immature naive state as a hallmark of JDM. Furthermore, we find that these changes in B cells are paralleled by signatures of Th2-mediated inflammation. Additionally, our work identified SIGLEC-1 expression in monocytes as a composite measure of heterogeneous type I interferon activity in disease. We applied network analysis to reveal that hyperactivation of the type I interferon response in all immune populations is coordinated with dysfunctional protein processing and regulation of cell death programming. This analysis separated the ubiquitously expressed type I interferon response into a central hub and revealed previously masked cell states. Together, these findings reveal the coordinated immune dysregulation underpinning JDM and provide novel insight into strategies for restoring balance in immune function.

## Intro

Juvenile Dermatomyositis (JDM) is part of a broad group of childhood-onset autoimmune conditions characterized by a type I interferon (IFN) gene signature and specific autoantibodies ranging from related systemic conditions like systemic lupus erythematosus (SLE) to endocrine-specific disorders like type I diabetes (1–4). Despite a shared IFN signature, JDM is associated with pathognomonic rashes and proximal muscle weakness resulting in distinct clinical phenotypes. The etiology of JDM is not fully understood but studies have shown that JDM is autoimmune-mediated and associated with a combination of genetic and environmental risk factors (5). While mortality is low with corticosteroid treatment, long-term patient follow-up studies have reported that 60-70% of patients have cumulative tissue damage (6, 7) with the risk of damage increasing almost linearly for each year after diagnosis (8). This finding highlights the importance of early disease intervention, the need to limit the harmful effects of long-term corticosteroid use in children (9), and the need for a personalized approach to disease management to improve upon these outcomes.

Clinical management of JDM currently relies on compiled empirical metrics such as physician global visual analog scale (VAS) of disease activity and muscle strength quantified via the childhood myositis assessment scale (CMAS) or manual muscle testing (MMT) (10). However, how these clinically observable phenotypes are rooted in disease immunopathology remains insufficiently understood. The presence of myositis-specific antibodies (MSA) that correspond to distinct clinical phenotypes and recent work showing that MSAs may be pathogenic (11,12) suggest the involvement of B cells (13). The expansion of naïve B cells in JDM has been highlighted by three independent studies using flow cytometry, mass cytometry, and single-cell RNA sequencing, respectively (14–16). The adaptive arm of the immune system is further implicated in disease pathogenesis by several large immunophenotyping studies that demonstrated the expansion of extra-follicular Th2 memory cells and central memory B cells (17,18). Additionally, the innate immune system has emerged as a contributor in JDM. Inflamed muscles of JDM patients exhibited the presence of plasmacytoid dendritic cells and macrophage-secreted proteins (19, 20), while similarly, biopsies of JDM and adult DM skin lesions showed an increase in CD14^+^ and CD68^+^ macrophages (21, 22). In peripheral blood, NK cells were found to be both dysfunctional and hyperproliferative in JDM (15, 23). Together, these findings highlight the involvement of both the adaptive and innate immune compartments in JDM across the peripheral immune system and disease-affected tissues. However, it also raises the question of whether the cause of JDM lies in a single cell type, or is a manifestation of broadly dysregulated cellular interactions across the immune system.

Systems-level studies based on single-cell measurements are required to reveal how dysregulated cell populations act individually or cooperatively to produce the observed chronic inflammation. Accordingly, several groups have turned to single-cell RNA sequencing as it enables unbiased profiling of tissues at single-cell resolution. In the first single-cell study of peripheral blood of JDM patients, we previously described a pan-cell-type IFN gene signature over-expressed in treatment-naive JDM that was most strongly correlated with disease activity in cytotoxic cell types (16). This signature has since been independently identified in the peripheral blood of treatment-naive patients (24). However, these studies have utilized small cohorts and lack adequate controls, in part due to the rarity of JDM in the human population. Thus, it has been challenging to determine which of these cell populations are specific to JDM, how these disease-specific dysregulated cell states are coordinated with one another, and which of these states cooperatively change in response to treatment.

In this study, we addressed this challenge by profiling JDM across several stages of disease activity using multiplexed Cellular Indexing of Transcriptomes and Epitopes by sequencing (CITEseq) of peripheral blood mononuclear cells (PBMCs) from 15 JDM patients, totaling 22 samples, and 5 healthy controls (HC). To minimize confounding by immune suppression, we included 9 treatment-naive samples as well as 6 samples from patients with inactive disease off medication. We leveraged CITEseq’s simultaneous profiling of gene and surface protein expression to gain more insight into cell phenotype and function, as cell surface proteins are used as diagnostic biomarkers and the targets of many biologic drugs. We first analyzed the data based on compositional differences in immune populations and confirmed the disease activity-associated imbalance of naive and mature lymphocytes. We next performed a multi-modal differential expression analysis and identified surface protein expression of SIGLEC-1 in CD14^+^ monocytes as a composite metric of the transcriptional Type I IFN response and clinical disease activity. To move beyond the identification of disease-specific cell populations and towards an understanding of immune-scale dysregulation in JDM, we applied a recently developed computational method DECIPHERseq (25) to infer networks of coordinated cell states from large cohorts of single-cell data. Importantly, this unsupervised network inference approach takes advantage of the biological heterogeneity in our entire dataset, improving upon previous work that relied on traditional pairwise comparisons of subsetted disease groups. Our analysis revealed immune dysregulation of cell homeostasis processes in active disease and imbalance of mature and naive lymphocytes expressing signatures suggestive of extra-follicular reactions as a potential autoimmune mechanism.

## Results

### JDM is associated with immunophenotypic differences in B and CD4+ T cell compartments

In order to gather a dataset with appropriate controls and limited confounding, patients were selected according to disease activity and medication status (Table 1). CITEseq was performed on PBMCs to generate single-cell libraries (Figure 1). Surface protein expression was measured using antibody-derived tags (ADT). Following pre-processing steps, we identified 29 clusters, which comprised 21 distinct immune cell populations across 105,827 cells. Clusters were annotated using canonical RNA and protein markers (Supplemental Figure 1A-B) within all major mononuclear immune cell compartments (Fig 2A).

**Table 1.** Disease Characteristics.

|  | TNJDM<br>(N=9) | Active<br>(N=7) | Inactive<br>(N=6) | HC<br>(N=5) |
| --- | --- | --- | --- | --- |
| **Sex** |  |  |  |  |
| Female | 3 (33.3%) | 4 (57.1%) | 5 (83.3%) | 2 (40.0%) |
| Male | 6 (66.7%) | 3 (42.9%) | 1 (16.7%) | 3 (60.0%) |
| **Age (years)** |  |  |  |  |
| Median [min-max] | 7.0 [2.0-15] | 13 [5.0-18] | 16 [5.0-18] | 3.0 [1.0-16] |
| **Prednisolone treatment — no. (%)** | 0 (0%) | 1 (14.3%) | 0 (0%) |  |
| **Methyl prednisolone treatment — no. (%)** | 0 (0%) | 1 (14.3%) | 0 (0%) |  |
| **Methotrexate treatment — no. (%)** | 0 (0%) | 4 (57.1%) | 0 (0%) |  |
| **HQL treatment — no. (%)** | 0 (0%) | 4 (57.1%) | 0 (0%) |  |
| **IVIG treatment — no. (%)** | 0 (0%) | 1 (14.3%) | 0 (0%) |  |
| **Physician Global VAS** |  |  |  |  |
| Median [min-max] | 5.0 [1.5-7.5] | 1.5 [0.50-8.0] | 0 [0-0.10] |  |
| **Muscle VAS** |  |  |  |  |
| Median [min-max] | 4.0 [0.50-8.5] | 0.50 [0-3.5] | 0 [0-0] |  |
| **Cutaneous VAS** |  |  |  |  |
| Median [min-max] | 4.0 [1.5-7.0] | 1.0 [0-7.0] | 0 [0-0.20] |  |
| **Patient/Parent Global VAS** |  |  |  |  |
| Median [min-max] | 5.0 [2.0-8.2] | 2.0 [0-8.0] | 0 [0-0.40] |  |
| Missing | 2 (22.2%) | 0 (0%) | 0 (0%) |  |
| **CDASIACT** |  |  |  |  |
| Median [min-max] | 16 [4.0-35] | 2.0 [0-8.0] | 0 [0-1.0] |  |
| **CHAQ-score** |  |  |  |  |
| Median [min-max] | 1.0 [0-2.4] | 0 [0-1.3] | 0 [0-0.38] |  |
| Missing | 2 (22.2%) | 0 (0%) | 0 (0%) |  |
| **MMT8-score** |  |  |  |  |
| Median [min-max] | 70 [59-76] | 80 [67-80] | 80 [79-80] |  |
| Missing | 4 (44.4%) | 0 (0%) | 0 (0%) |  |
| **MSA** |  |  |  |  |
| MDA5 | 1 (11.1%) | 0 (0%) | 0 (0%) |  |
| NEG | 1 (11.1%) | 1 (14.3%) | 1 (16.7%) |  |
| NXP2 | 1 (11.1%) | 4 (57.1%) | 2 (33.3%) |  |
| TIF1y | 6 (66.7%) | 2 (28.6%) | 2 (33.3%) |  |
| UNK | 0 (0%) | 0 (0%) | 1 (16.7%) |  |
| **Muscle enzyme elevation Present — no. (%)** | 8 (88.9%) | 3 (42.9%) | 1 (16.7%) |  |

Overview of clinical characteristics from the 15 patients with JDM and 5 healthy controls, totalling 27 samples. Some patients had longitudinal sampling at different disease stages. Myositis specific antibodies were sent to Oklahoma Myositis Research Foundation for testing. TNJDM = treatment-naïve JDM, Active = Active JDM defined by Physician global visual analog score (VAS) $\geq 0.5$ and on medication, Inactive = Inactive JDM defined by physician global VAS $< 0.5$ and off medication, HC = healthy control, HQL = Hydroxychloroquine, VAS = Visual Analogue Scale, CDASIACT = Cutaneous Dermatomyositis Disease Area and Severity Index Activity Score, CHAQ = Childhood Health Assessment Questionnaire, MMT = Manual Muscle Testing, MSA = Myositis Specific Antibodies.

**Figure 1:**
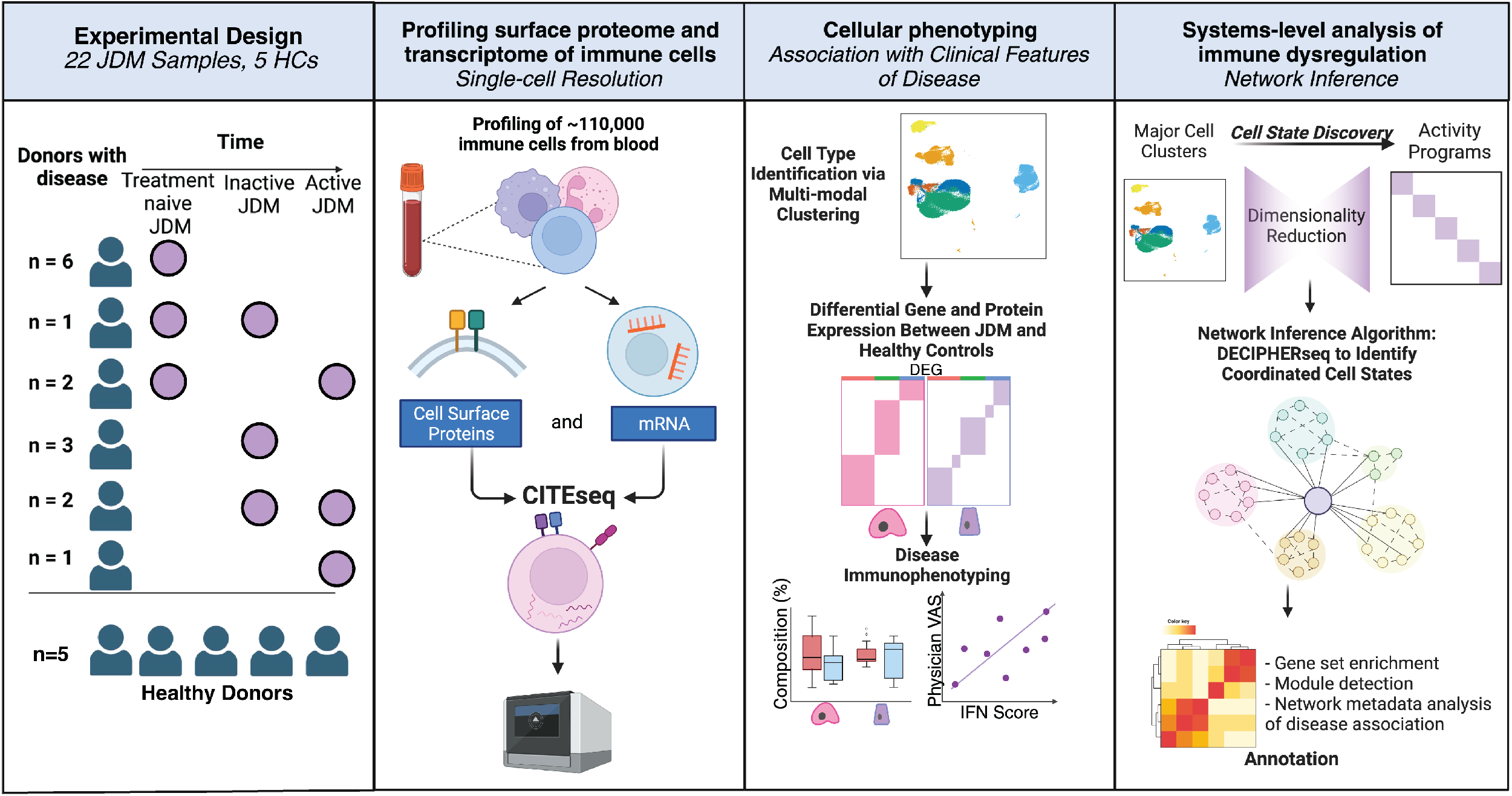
Study design and analysis strategy for profiling PBMCs from 27 samples (n=22 JDM, n=S HC)

**Figure 2.**
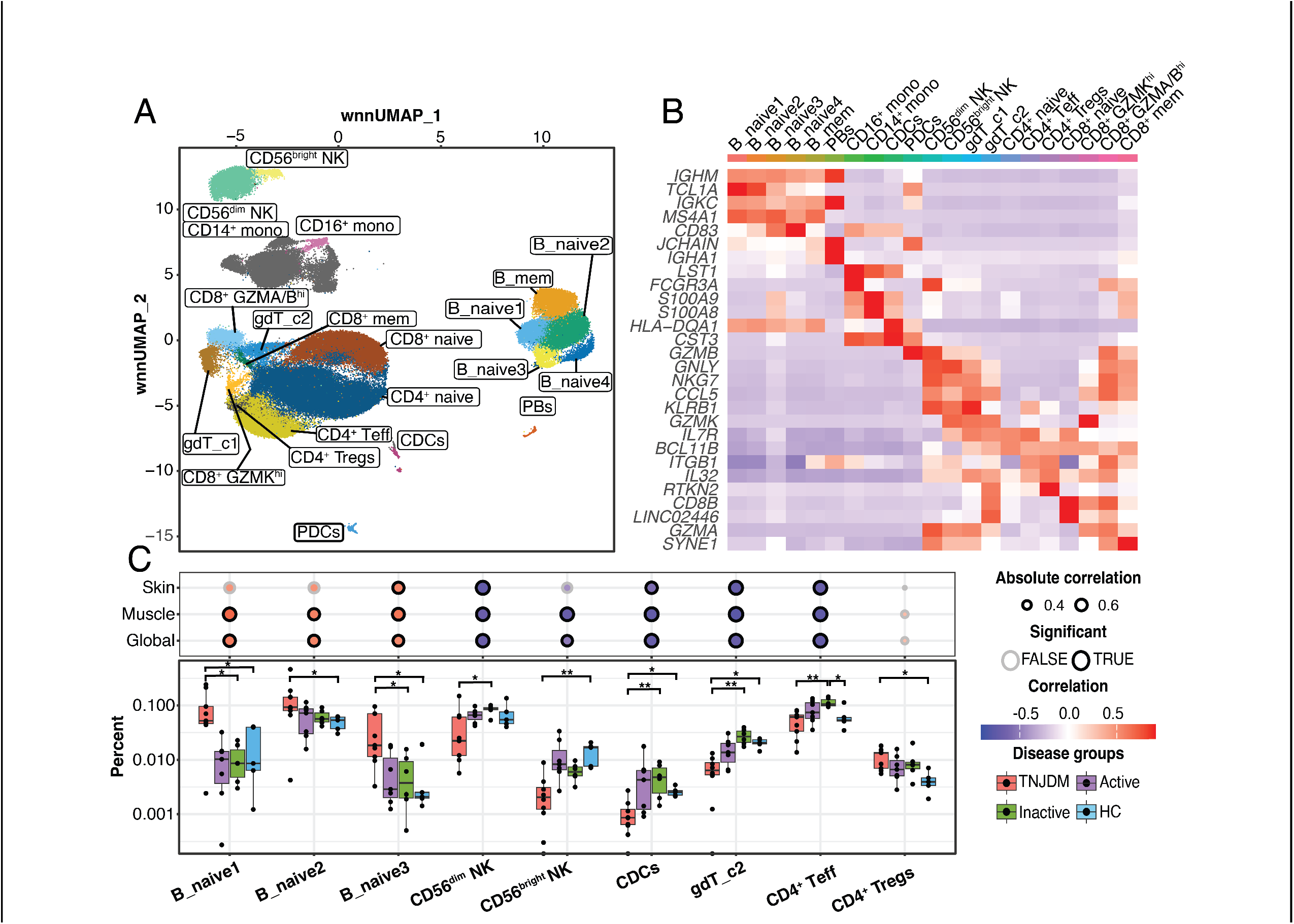
Cell types associated with JDM in peripheral blood. **(A)** UMAP constructed using weighted-nearest neighbors (wnn) clustering colored by cell type. pDCs=plasmacytoid dendritic cells, cDCs=classical dendritic cells, PBs=plasmablasts. B_mem = memory B cells. **(B)** Heatmap with top 2 markers per cluster. **(C)** Boxplot shows cell type proportion by disease group, using Kruskal-Wallis test with Dunn’s post hoc test comparing TNJDM to HC, TNJDM to inactive JDM and inactive to HC (Holm p.adj < 0.05). The dotplot above shows the Spearman correlation between corresponding cell type proportion in boxplot and Physician Global VAS, where the size of the dot indicates the correlation, the color indicates the direction of the correlation, and the border weight indicates significance (p.adj < 0.05).

We first characterized global changes to cell composition across disease states comparing treatment-naive JDM, inactive JDM and HC. We quantified the differences in cell type proportion between disease states and the correlation to disease activity measures for physician global (PGA), cutaneous, and muscle VAS scores. Within the T cell compartment, the proportion of regulatory T cells (Tregs) (CD45RO^+^, IL2R^+^, *FOXP3^+^*) was increased in patients with treatment-naive JDM (p = 0.02) consistent with previous findings (16). CD4^+^ effector T cells (CD45RO^+^) and gdT cluster 2 (*TRDC,TRGC*) were significantly increased in patients with inactive JDM and the proportion of cells from these populations negatively correlated with disease activity measures (p ≤ 0.05, Spearman). There was an overall decrease in innate populations, including CD56^bright^ and CD56^dim^ NK cells and classical dendritic cells (cDCs) in treatment-naive JDM compared to HC and inactive JDM, and the proportion of these cell types also correlated negatively with disease activity (p < 0.05, Spearman).

Compared to healthy controls and patients with inactive disease, treatment-naive patients had higher proportions of multiple naive B cell populations, including B_naive1 (IgM^+^IgD^+^CD38^+^CD24^+^CD10^+^) corresponding to an immature naive B population, B_naive2 (IgM,^+^IgD^+^CD38^lo^CD24^lo^), and B_naive3 (IgM^+^IgD^+^CD38^+^CD24^+^), and the proportion of these populations positively correlated with multiple disease activity measures (p < 0.05, Spearman) (Fig 2C). The proportion of B_mem cells, characterized by *TNFRSF13B* (encoding TACI) expression, negatively correlated with the muscle VAS score (p < 0.05, Spearman). B_naive4 consisted solely of cells from one treatment-naïve patient and one healthy control and was excluded from the analysis, and clustering was weighted highly for RNA, suggesting a distinct gene signature was driving this separation. An orthogonal analysis clustering all B cells by ADT alone resolved patient-specific clustering and confirmed expansion of the immature naive B population (IgM^+^IgD^+^CD38^+^CD24^+^CD10^+^) and relative decrease of memory B populations in treatment-naive JDM highlighting skewing of the B cell compartment toward an immature state at disease onset (Supplemental Figure 2). The immature naive B population in both analyses had higher expression of CD38 (both RNA and protein) and *MZB1,* two genes essential for plasma cell differentiation, than all other B cell clusters (26, 27).

Given the observed imbalance of lymphocytes in treatment-naive JDM, we next sought to immunophenotype B cell and CD4^+^ T cell subsets in JDM at the proteomic level to gain molecular insight into cell states. Differential protein analysis of immature naive B cells comparing treatment-naive JDM to HC identified increased expression of CD325 and MICA-MICB and decreased expression of CD1C, BAFF-R and PD-L1 (Supplemental Table 1). Within the CD4^+^ T compartment (Supplemental Figure 3A), CD4^+^Tregs from TNJDM had higher expression of Tim-3, ICOS, CD164 and CD38 and down-regulation of CD101, a molecule which decreases pro-inflammatory T cell responses (28). CD4^+^Teff in patients with treatment-naive JDM had higher surface expression of CD164 and PD-1 and down regulation of KLRG1, an inhibitory molecule (Supplemental Figure 3B). The over-expression of PD-1 on the cell surface suggested that peripheral T helper cells might be present in JDM (29, 30). However, while ICOS expression was higher (Benjamini-Hochberg (BH) p < 0.05), no difference was found in surface expression of CXCR5 between CD45RO^hi^PD-1^hi^CD4^+^ T cells and CD45RO^lo^PD-1^lo^CD4^+^ T cells, and these cells were not significantly expanded in JDM (Supplemental Figure 3C-E). Taken together, these compositional and immunophenotyping observations add to the growing body of work showing that JDM in the treatment-naive state is characterized by relative imbalances of naive and mature lymphocyte states (16, 24), reduced innate immune populations (23) and distinct CD4^+^ T and B cell immunophenotypes (17,18).

### SIGLEC-1 expression is a composite measure of the IFN gene signature in JDM

We next compared gene and protein expression between treatment-naive JDM and HC samples in all cell types based on the hypothesis that certain cell types may not be altered in composition but may be functionally altered at the molecular level. Monocytes displayed the highest number of differentially expressed genes (n = 211) and proteins (n = 19) in our pairwise analysis including CD169 (SIGLEC-1), CD107a (LAMP-1), and CD164 (Supplemental Figure 4). SIGLEC-1 is a monocyte-restricted IFN-induced protein that was recently identified as a potential biomarker in JDM (31). Both CD107a and CD164 are cell adhesion molecules involved in trafficking of activated mononuclear cells and adhesion to vascular endothelium (32).

A common finding across all cell types was overexpression of genes enriched in Type I IFN processes, which was previously reported in bulk expression data (33, 34) and confirmed in single-cell studies (Supplemental Figure 5) (16, 24). Taking advantage of our richer dataset, we also identified that the strength of the IFN gene signature varied widely across both patients (Fig 3c) and cell types. Using an IFN gene score derived from the transcriptional data (Supplemental Figure 6), we plotted the average score per patient per cell type and applied hierarchical clustering, which identified a cluster of “IFN-hi” patients and “IFN-lo” patients. The group of “IFN-lo” patients included two treatment-naive JDM patients that clustered with inactive and HC patients. This heterogeneity of the IFN gene signature was, in part, explained by disease activity level (Fig 3d) as a bulk IFN gene score significantly correlated with disease activity (R=0.69, Spearman), but also highlights that some patients with JDM have little to no IFN gene expression detectable in PBMCs suggesting heterogeneity of this gene signature in a subset of patients with JDM.

**Figure 3.**
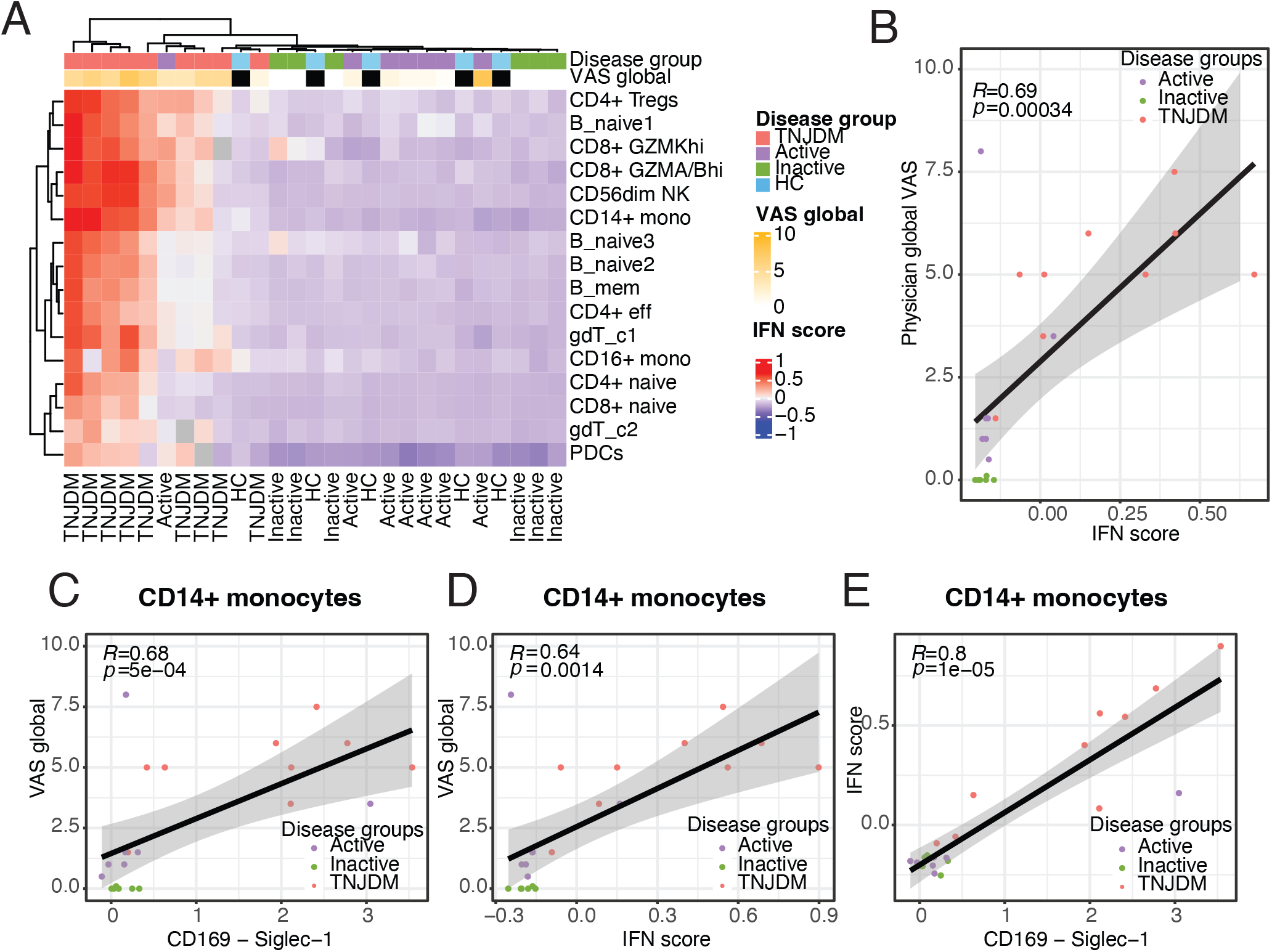
Type I IFN-induced gene and protein expression is associated with disease activity in JDM. **(A)** Heatmap of average IFN score per cell type and sample. Hierarchical clustering was performed using Euclidean distance and the complete clustering method. IFN score was calculated based on average expression of IFN across all cells per sample. **(B)** Spearman correlation between IFN score and Physician Global VAS colored by disease group.**(C)** Scatter plot showing Spearman correlation between CD169 (SIGLEC-1) expression and IFN score for CD14+ monocytes.**(D)** Scatter plot showing Spearman correlation between CD169 (SIGLEC-1) expression and Physician Global VAS for CD14+ monocytes. **(E)** Scatter plot showing Spearman correlation between IFN-score and Physician Global VAS for CD14+ monocytes.

Given that SIGLEC-1 is a type I IFN-induced protein, we next wanted to determine if patterns of type I IFN stimulated gene expression were reflected at the protein level, as protein biomarkers are more amenable for clinical lab-based testing. SIGLEC-1 expression in CD14^+^ monocytes correlated with disease activity to a similar degree as the IFN gene signature (Fig 3e and 3f), and SIGLEC-1 expression was itself highly correlated with the IFN gene signature in monocytes (Fig 3g). This suggests that SIGLEC-1 expression in CD14^+^ monocytes is a representative composite measure of the IFN gene signature in JDM and supports the role of type I IFN in the immunopathology of JDM. Overall, these results underscore the potential of SIGLEC-1 as a biomarker of IFN response in JDM that may be useful for stratifying disease severity and tracking disease activity.

### Unsupervised network analysis reveals coordinated cell states shared among immune cells in JDM

Given that differential expression analysis relies on pairwise comparisons between only subsets of the data, we next aimed to identify biological features of JDM that are shared across stages of disease activity. We therefore applied an unsupervised network inference method, DECIPHERseq, to the 6 major cell types annotated in the dataset: B cells, CD4T, CD8T, NK cells, gdT cells, and myeloid cells. These cell populations were chosen to pass thresholds of at least 100 cells of a given cell type in each sample. DECIPHERseq relies on non-negative matrix factorization (NMF) (35–37) to first break the dataset down into gene sets that represent distinct states of biological activity, or ‘activity programs’, and then constructs a network of gene expression programs (GEPs) based on how expression of the programs covaries across patient samples (Figure 4A). After outlier filtering, NMF identified 76 activity programs (Figure 4B).

**Figure 4.**
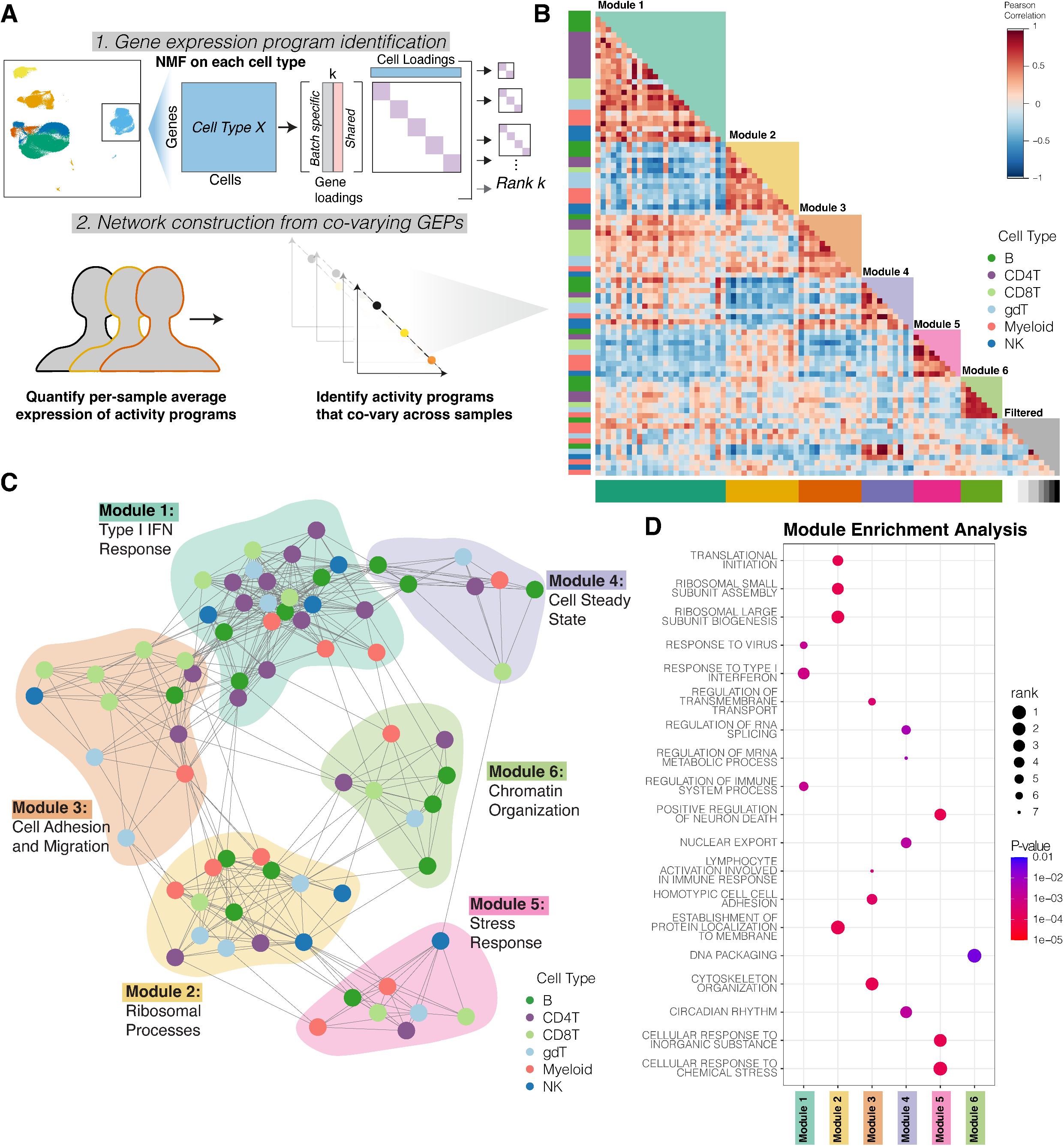
DECIPHERseq reveals network of coordinated GEPS from scRNAseq data in JDM. **(A)** Overview of the OECIPHERseq workflow. **(B)** Heatmap showing 6 major clusters of GEPs identified by OECIPHERseq (Pearson). GEPs are clustered into modules, with isolated GEPs filtered out (greyscale). **(C)** Force-directed network constructed from correlated GEPs in PBMCs from JOM patients and healthy controls. Nodes represent programs in the given cell types and edges represent positive significant correlations (Pearson, p<0.05). **(D)** Ootplot showing selected gene sets found to be enriched within specific modules compared to the rest of the network. Color corresponds to module enrichment p value and size corresponds to a set’s rank in list of significantly enriched gene sets for that given module ordered by ascending module enrichment p-value. All gene sets shown fall in the top 10 terms for their respective modules (total gene sets: 626).

Next, a force-directed network graph from the correlation matrix of activity programs was constructed where each node represents a program, and each edge represents a statistically significant positive correlation between two nodes. Correlations between programs are accounted for in the network visualization such that further apart nodes can be interpreted as being negatively correlated and closer nodes can be interpreted as being positively correlated programs (Figure 4C). Using DECIPHERseq’s community detection algorithm, we identified 6 hubs of inter-connected activity programs or ‘modules.’ All modules contained multiple cell types, highlighting that many biological processes in JDM are coordinated across several immune cell types. We annotated each node using gene set enrichment analysis (38, 39) of gene ontology terms (GO) (40) on each program’s ranked gene list (Supplemental Table 2, Supplemental Figures 7-12).

DECIPHERseq’s module enrichment analysis identified consensus biological themes for each module in an unsupervised manner by quantifying the likelihood of a GO term being shared across the programs in a module if programs were randomly sampled from the entire network by chance (Figure 4D, Supplemental Table 3). Module 1 was enriched for Type I IFN response programs including gene sets such as ‘Response to Virus’ and ‘Response to Type I IFN’. Module 2 consisted of programs enriched for protein assembly genes used in ribosomal processes including ‘Translational Initiation’ and ‘Ribosomal Large Subunit Biogenesis’. Module 3 was comprised of mostly lymphocyte programs with strong correlations to Type I IFN response programs in Module 1. Module 3 was significantly enriched for gene sets related to cell adhesion and migration suggesting that this module identified programs that represent lymphocyte extravasation to tissue. Module 4 represented cells’ steady state processes as it was enriched for gene sets like ‘Circadian Rhythm’. Module 5 was annotated as a Stress Response module because it was enriched for ‘Cellular Response to Inorganic Substance’ and ‘Cellular Response to Chemical Stress’. Module 6 contained very few gene sets that were unique to the module, as it consisted of programs enriched for programs intrinsic to eukaryotic cells like ‘DNA Packaging’ and ‘Chromatin Organization.’

### JDM CD4T cells express Th-2 programming coordinated with B cell and IFN responses

Next, we aimed to interpret the annotated network in the context of GEPs associated with JDM compared to healthy control patients irrespective of disease activity. In the annotated network, we first focused on Module 1, which was enriched in type I IFN responses and many programs in this module were increased in TN-JDM, as expected (Figure 5A). All 6 major cell types expressed an IFN gene program which were highly correlated to one another, as shown by the closely connected hub at the center of module 1. This IFN hub was associated with JDM as compared to HC patients (t-test, p<0.05). Using DECIPHER-seq, we identified additional coordinated gene programs in Modules 1-3 expressed more highly in all JDM patients compared to HCs, irrespective of disease activity. This is a strength of DECIPHER-seq, which uses NMF to uncover co-occurring activity programs that were previously obscured in DGE analysis by the high number of overexpressed IFN-related genes.

**Figure 5.**
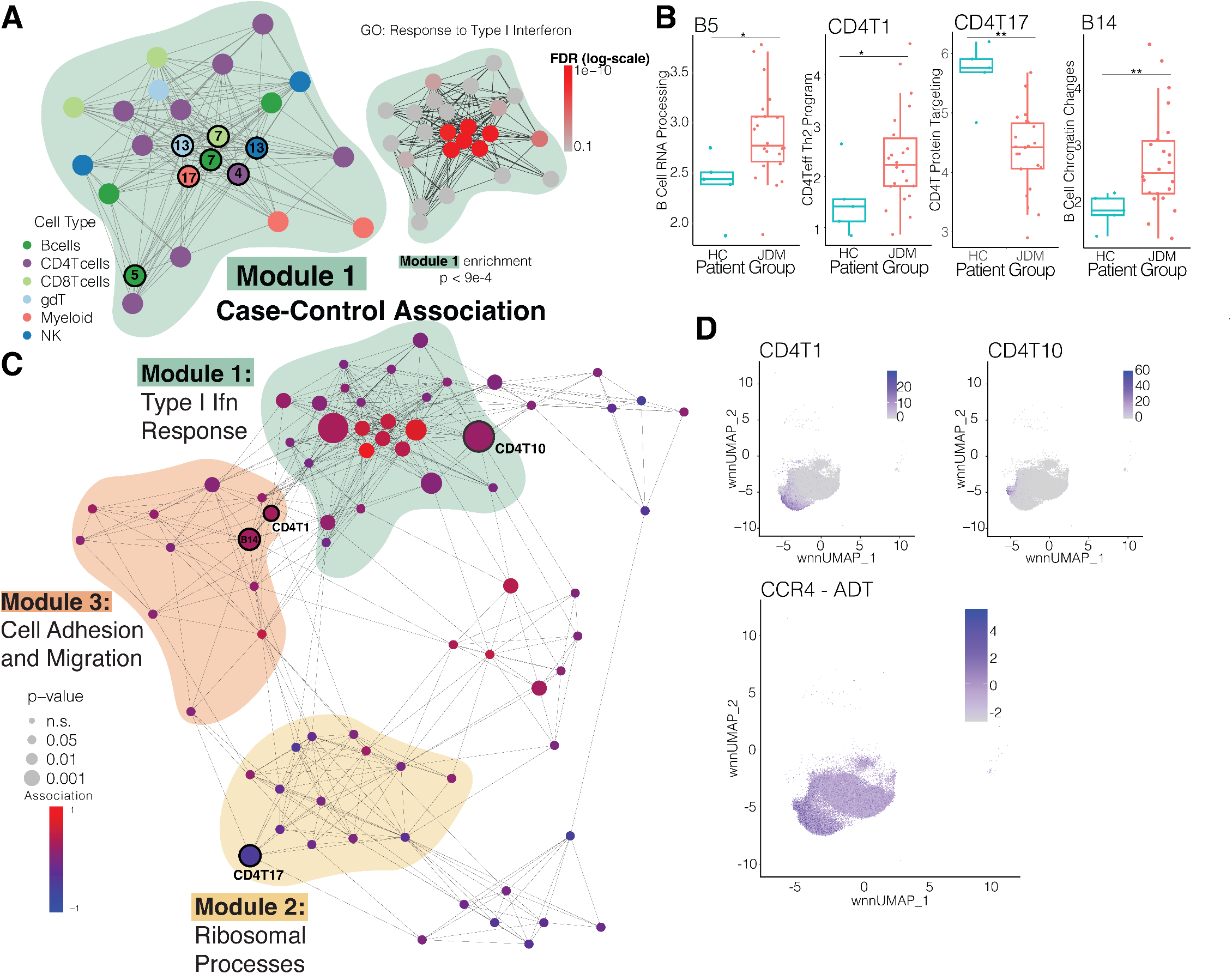
JDM is associated with a central IFN hub and cell specific gene programs in the B and CD4T compartments. **(A)** Zoomed in graph of Module 1. GSEA results for Response to Type I IFN GO term shown with each node colored according to FDR. Adjusted p-value of module enrichment is also shown. **(B)** Boxplots showing significant differences in expression of selected programs between HG (n=5) and JDM patients (n=22). (* p<0.05, ** p<0.01) (C)Network graph showing case-control analysis of each program’s expression, with node size scaled according to p-value and colored according to strength of the association between disease status and program expression (t-test). **(D)** wnnUMAPs showing expression of GEPs CD4T1 and CD4T10 with co-expression of surface protein CCR4.

Reflecting the altered composition of lymphocytes in JDM, several lymphocyte programs were significantly associated with JDM (p<0.05). In addition to IFN gene programs, JDM patients more highly expressed two B cell programs: B5 in Module 1 (IFN Response) was enriched in mRNA metabolic processing, RNA splicing, chromatin organization and modification and cell cycle and B14 in Module 3 (Cell Adhesion and Migration) was enriched in chromatin remodeling and cytoskeletal organization (Supplemental Figures 7, 9). These enriched biological processes suggest that a subpopulation of B cells are more transcriptionally active and undergoing epigenetic regulation in JDM relative to healthy controls. In Module 3, correlated to B14, CD4T1 (enriched in cell migration, adhesion, activation, and secretion) was expressed more highly in JDM and in the region of the UMAP corresponding to CD4^+^Teff cells. This CD4T1 program expressed by CD4^+^Teff cells contained several genes (*GATA3*, *CCR4*, *PRDM1*) that indicate skewing towards a Th2 subset while expression of *PRDM1* (Blimp-1) suggests participation in extra-follicular reactions. Th2 CD4^+^ T cells were previously found to be expanded in JDM and associated with extra-follicular B-T cell help (17, 18). We observed similar expression of Th2 genes (*GATA3*, *CCR4*, *PRDM1*) in CD4T10, a Treg program (*FOXP3*, *IKZF2*, *IL2RA*) expressed more highly in JDM. Notably, CD4T17 was negatively associated with JDM; this program was enriched in protein targeting to the membrane and endoplasmic reticulum indicating that these processes may be dysfunctional in CD4T cells of children with JDM. Together, these results suggest that though CD4^+^T cells are skewed towards a Th2 phenotype with the capacity to help B cells via extra-follicular B-T cell interactions, protein processing of these cells may be dysfunctional. DECIPHERseq revealed that this Th2 alteration in the CD4^+^ Treg and Teff cells is accompanied by the emergence of a transcriptionally active subpopulation of B cells, suggesting coordinated alterations in JDM lymphocytes.

### Novel cell states are correlated with IFN gene expression in treatment-naive JDM

We next wanted to identify modules and gene programs associated with stages of disease activity in JDM (HC, Inactive and Active JDM, and treatment-naive JDM). To do so, we performed a 4 group ANOVA on each program in the network and post-hoc pairwise analysis using the Tukey test (Figure 6A). We identified programs in Module 1, 2 and 5 that were significantly associated with disease activity (ANOVA p<0.05). The IFN gene programs were also significantly overexpressed in treatment-naive JDM patients, as expected. Notably, expression of the central Module 1 IFN hub GEPs in all six major cell types more strongly correlated to the clinically evaluated PGA than the pseudobulk IFN gene score derived from pairwise DEG analysis (Supplemental Figure 13), underscoring the utility of an integrative approach like DECIPHERseq in uncovering clinically relevant gene signatures.

**Figure 6.**
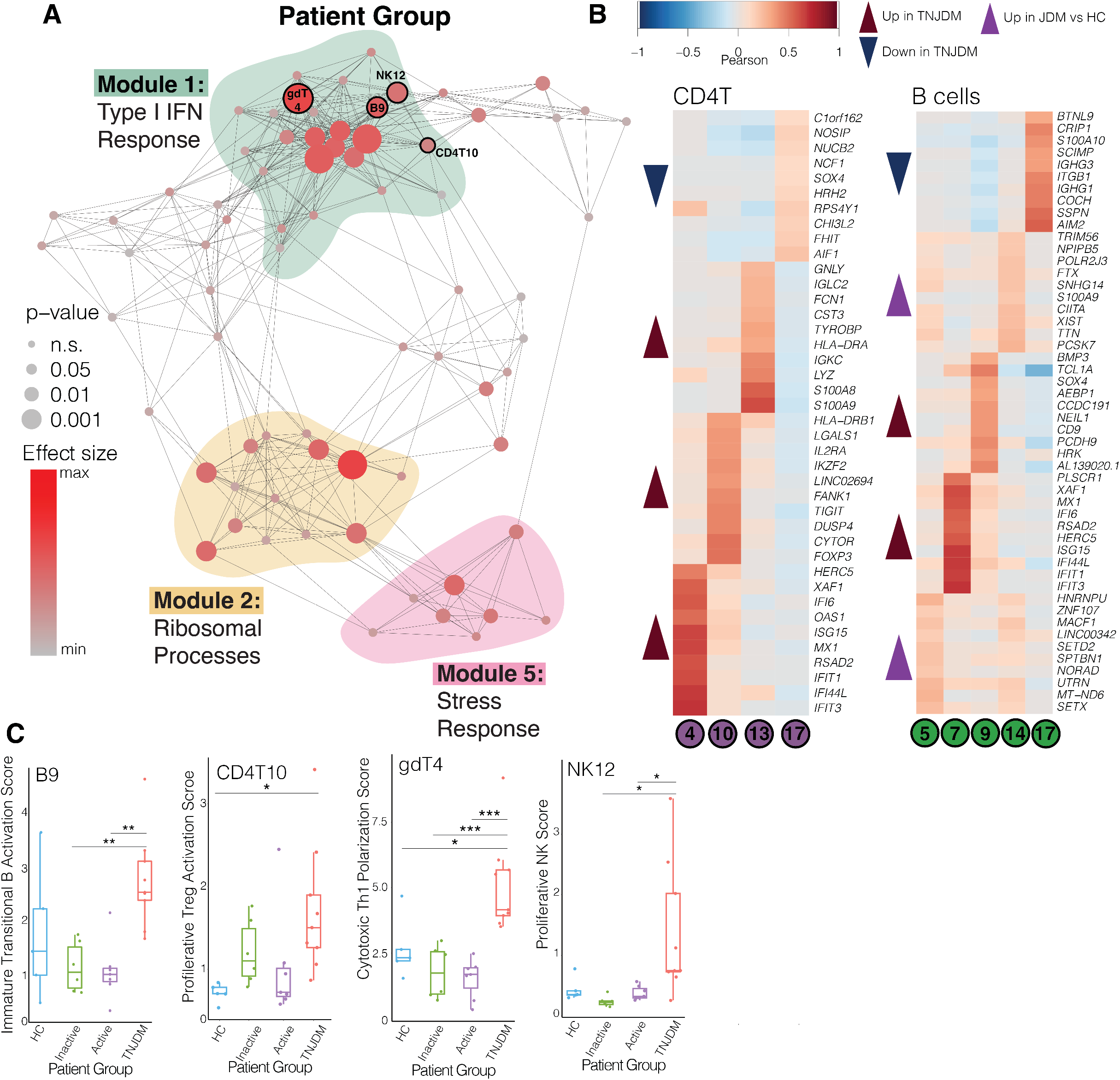
Disease activity in JDM is associated with central hub of IFN response in network, correlated with novel cell states. **(A)** Network graph showing results of 4-group ANOVA of each program’s expression, with node size scaled according to p-value and colored according to strength of the association between disease status and program expression. **(B)** Heatmaps showing top 1O marker genes for selected disease-associated programs for CD4T and B cells. Colored according to Pearson correlation between gene expression and program expression in the indicated cell type. **(C)** Boxplots showing significant differences in expression of selected programs between HC (n=5), Inactive JDM (n=6), Active JDM (n=7), and TNJDM patients (n=9). (* p<0.05, ** p<0.01, ***p<0.001).

By isolating these IFN GEPs in each cell type, we were able to determine disease activity-associated programs correlated with the IFN hub, some of which corroborate previous findings (Figure 6A, C). We identified B9, an immature naive B cell program (*CD24*, *CD38*, *MME*)), to be significantly associated with disease activity (Figure 6B). This gene program shared several top markers (*TCL1A, SOX4, NEIL1*) with the immature B cell population that we and others previously found to be expanded in treatment-naive JDM (16, 24). Notably, expression of this activated immature B cell program could be attributed to the B_naive1 cluster that we observed to be increased in treatment-naive JDM during our previous compositional analysis (Supplemental Figure 14). Similarly correlated with the IFN hub, NK12 was associated with treatment-naive JDM compared to active and inactive disease (p<0.05, Tukey). NK12 (*MKI67, HIST1H1B*) was enriched for gene sets related to cell proliferation and epigenetic regulation, confirming findings that a subset of NK cells in JDM are highly activated and proliferative (Figure 6C) (15, 23).

Given our ability to identify cell states shown in prior work to be relevant in JDM with this approach, we next focused our attention on the novel disease activity-associated programs that DECIPHERseq identified as correlated with the Module 1 IFN hub. We identified CD4T10—significantly associated with disease relative to HCs in both treatment-naive JDM and all JDM cases suggesting a central role in JDM (Figures 6B, 7C)—as a proliferative Treg program which includes signatures of activated Th2 effector cells implicated in extra-follicular B-T interactions (*FOXP3*, *IL2RA*, *PRDM1*, *MKI67, GATA3, CCR4*) (41). CD4T10 also was characterized by high expression of *IKZF2*, an important transcription factor for thymocyte development, indicating that cells that express CD4T10 are thymically generated Tregs (tTreg) as opposed to peripherally induced Tregs (42–44). Notably, CD4T10 included the marker *CCR4*, a chemokine receptor highly expressed in Tregs that are preferentially recruited to skin under inflammatory conditions (45). Expression of CD4T10 co-localized with surface protein expression of CCR4 in the UMAP as well (Figure 5D), highlighting the advantage of this multi-modal sequencing approach in identifying functional markers of transcriptomic signatures.

We also identified the program gdT4, a cytotoxic Th1 polarized gdT program (*GZMB, CX3CR1, TBX21*) that was also correlated with the central IFN hub and was significantly increased in treatment-naive patients compared to both active and inactive JDM and HC (p<0.05, Tukey). High expression of *TRGC1* and *TBX21*, encoding the transcription factor T-bet responsible for regulating IFNG expression, specifically identified cells expressing this program as Th1-like TCRVd1 gdT cells (46, 47). A similar subpopulation of gdT cells was found to be increased in synovial fluid and blood of juvenile idiopathic arthritis patients, another disease thought to be mediated by autoantibody-driven recruitment of autoreactive immune cells to tissue (48, 49). This points to gdT4 as an important inflammatory cell state specific to treatment-naive disease that is potentially up-or downstream of the Type I IFN response broadly upregulated across immune cell populations in JDM. Together, CD4T10 and gdT4 highlight the utility of dimensionality reduction-based methods like DECIPHERseq in furthering our ability to interpret novel cell states in noisy high-dimensional data.

### Cell homeostasis processes are dysregulated across many immune cell types in JDM

All of the novel disease activity programs that were highly expressed in treatment-naive JDM were components of Module 1 which was enriched for Type I IFN and its associated immune processes. The network-wide ANOVA analysis also revealed disease activity-associated programs in Module 2 and Module 5, which were significantly anti-correlated with Module 1 (Figure 7A) and expressed at lower levels in treatment-naive JDM patients compared to healthy controls and other JDM patients. Module 2 was significantly enriched for gene ontology terms like ‘ribosome assembly’ and ‘translational initiation’ while Module 5 was enriched for terms like ‘regulation of cell death’ and ‘cellular response to chemical stress’ (module enrichment p<0.005) (Figure 7B). The disease-associated programs within these modules were expressed significantly lower in treatment-naive JDM, suggesting dysfunction of cellular processes that underpin ribosomal activity and cell stress response in at disease onset (Figure 7D).

**Figure 7.**
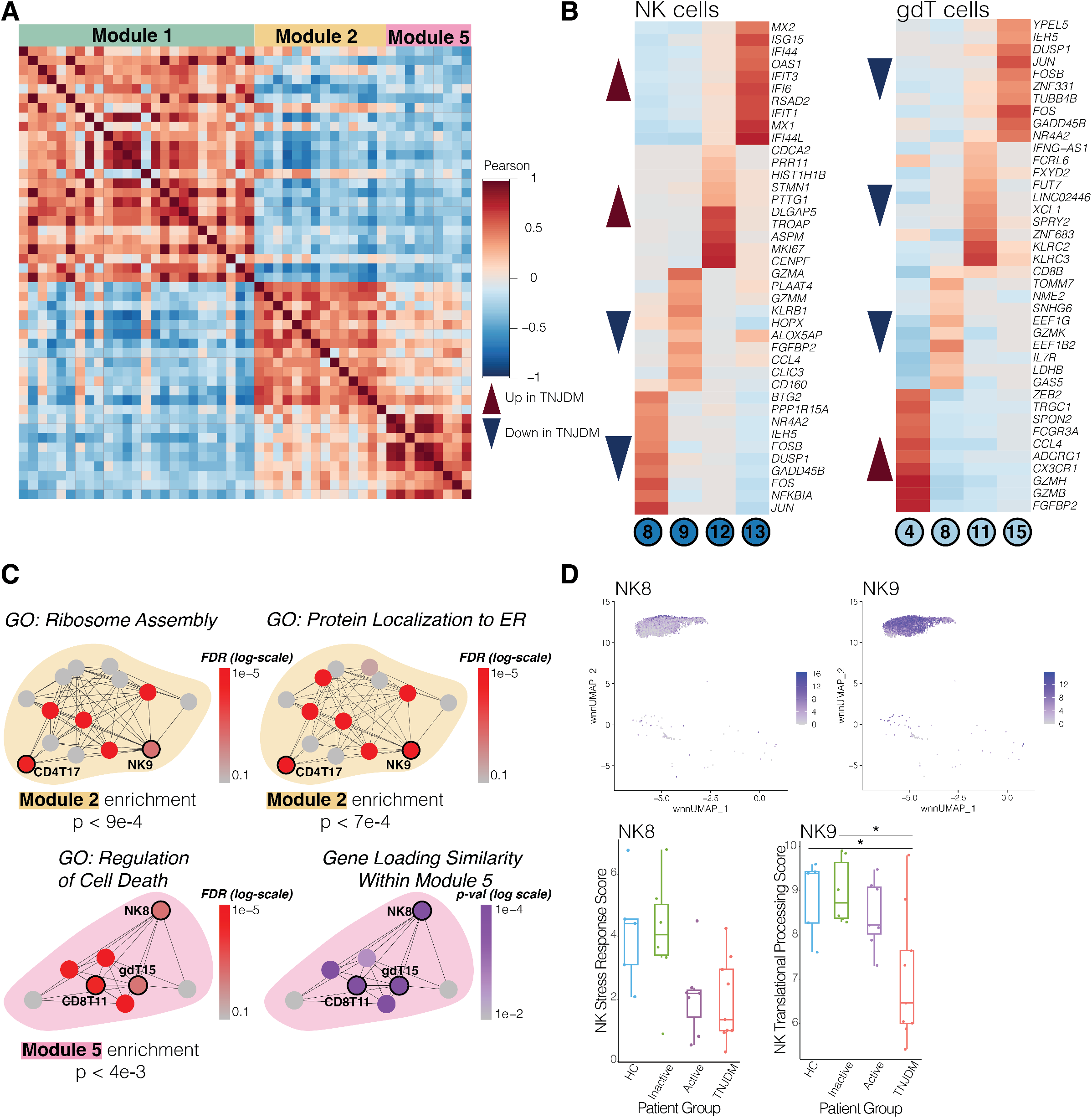
Protein processing and regulatory cell death programs are dysregulated in JDM. **(A)** Subset of Modules 1, 2, and 5 from original heatmap in Figure 4B highlighting the negative correlations. **(B)** Heatmaps showing top 10 marker genes for selected disease-associated programs for NK and gdT cells. Colored according to Pearson correlation between gene expression and program expression in the indicated cell type (p<0.05). C) Selected network modules are colored by FDR of enrichment for indicated gene ontology set ((FDR<0.01) or gene loading similarity within Module 5. D) wnnUMAPs showing single-cell expression of NK programs 8 and 9, and boxplots showing per-sample expression (4-way ANOVA p<0.05, Tukey *p<0.05).

Notably, disease activity-associated programs CD8T11, NK8 and gdT15 (*FOS, JUN, NFKBIA, CXCR4, SOCS1*) in Module 5 share their top 5 expressed genes (Figure 7C) and are each individually enriched in ‘regulation of cell death’ and ‘regulation of cell cycle’ (Supplemental Figure 11). We quantified the overlap in gene expression between activity programs by Fisher’s exact test and confirmed the high gene loading similarity between programs in Stress Response Module 5 (Figure 7C). All three of these programs are expressed at lower levels in active and treatment-naive JDM and negatively correlated with activated cytotoxic cell signatures identified in Module 1 (Supplemental Figure 15). In addition to these three disease activity-associated programs, our gene loading similarity analysis revealed that the programs CD4T9 and B10 also share top marker genes (*FOS*, *JUN*, *NFKBIA*). This suggests that regulatory mechanisms of cell death are disrupted in cytotoxic cell populations in patients with active disease and these processes are also disrupted in lymphocytes.

In the NK cell compartment, Program 9 was characterized by *CD160*, an important regulatory molecule on NK cells, *CCL4*, and *CX3CR1* which is implicated in the recruitment of NK cells to inflamed tissues (50). Notably, processes enriched in NK9 such as ribosome assembly and protein targeting to the ER were also found to be downregulated in treatment-naive JDM (23). Program 8 and 9 were broadly expressed programs in the NK cells (Figure 7D) and were expressed lower in treatment-naive JDM, suggesting dysfunctional cell signaling in NK cells may play a role in JDM as other studies have shown (15).

In Module 2, CD4T17 was enriched in several gene sets related to protein processing such as protein targeting to the ER. Interestingly, this program was also characterized by high expression of several genes encoding members of the actin protein family (*ACTB*, *ACTG1*). Given the crucial role actin filaments play in antigen recognition during the formation of the immune synapse, dysfunction in components of that protein machinery could have significant effects on the immune system. Among other disease activity-associated programs, differential expression of CD4T17 between HCs and JDM patients persisted even in inactive disease. Taken together, disease activity-associated programs in Modules 2 and 5 highlight shared cellular processes that may be under-active in JDM, providing new insights into potential cellular mechanisms that accompany the known signature of overactive IFN-response in JDM.

## Discussion

Multiple components of the adaptive and innate immune compartments have been implicated in the pathogenesis of JDM. However, previous studies have been unable to uncover how multiple disease-associated cell states are coordinated to produce the observed autoimmune signature. Here in the largest single-cell study of JDM to date, we provide an unbiased, comprehensive picture of immune dysregulation. We first show that immune dysregulation in JDM manifests at the level of compositional imbalance of immune populations and that these compositional changes are correlated to clinical metrics of disease activity. Next, we identify distinct disease-associated molecular signatures of lymphocyte and myeloid subsets through differential analysis and demonstrate that of these markers, surface expression of SIGLEC-1 in CD14+ monocytes is a composite metric of disease activity and reflects the heterogeneous type I IFN response in disease (34, 51–53). Finally, using DECIPHERseq to integrate disease-associated programs, we uncover how this type I IFN response is coordinated with known signatures and novel cell states, generating new hypotheses for disease etiology.

Broadly, we observe that the composition of PBMCs changes in JDM, with an under-representation of innate immune cells and an expansion of lymphocytes with bias toward immature naive populations over memory phenotypes in B and CD4T cells. Within the B cell compartment, the distinctive transcriptomic and proteomic signature of immature naive B cells is consistent with what we and others previously reported in treatment-naive disease (16, 24). Given that autoantibodies are thought to play a role in disease pathogenesis, this skewing of the B cell compartment would seem counterintuitive. However, given recent findings emphasizing the importance of extra-follicular B cell differentiation pathways through which autoreactive “activated naive” B cells are precursors to antibody-secreting cells, we hypothesize that this skewing may be suggestive of extra-follicular reactions in JDM (29, 30, 54). In fact, the expanded immature naive population had higher expression of CD38 and MZB1, genes important for plasma cell differentiation. While these data indicate that naive B cell populations expanded are in JDM, the overall low expression of CD27 and CXCR5 across all B cells made it difficult for to conclude that this population matches the double negative B cell population associated with SLE. However, recent immunophenotyping work in a large cohort of JDM patients that found simultaneous expansion of CXCR5-central memory B cells and Th2 cells provides support for further investigation into extra-follicular B-T cell help in JDM (17).

Accompanying these imbalances in B cells, we observe complementary dysregulation in the T cell compartment that lends further support to the hypothesis of extra-follicular interactions in JDM. In populations of peripheral blood *FOXP3^+^* Tregs and CD4^+^ effector T cells, we identify a common signature of Th2 activation expressing genes associated with extra-follicular T cell responses (*GATA3, CCR4, PRDM1*) that is associated with JDM compared to HC, and treatment-naive JDM when disease activity is considered. These findings are consistent with previous work, which identified skewing of CD4^+^ T cells toward a Th2 phenotype in JDM and showed in vitro that peripheral Th2 cells were efficient in helping B cells, including stimulating antibody production (18). Other studies have reported that Tregs are expanded in muscle while peripheral blood percentages remain unchanged and that these Tregs in JDM have diminished suppressive capacity, raising the possibility of these cells being exhausted in disease (55). In contrast, our results show that the expanded population of peripheral Tregs are proliferative and activated (*MKI67*, *IL2RA*, *IRF4*), taking on a Th2 signature.

One possible explanation for these seemingly discordant findings is that Tregs in JDM may coopt the transcriptional machinery of effector cells for Th2-specific suppression, which was proposed by Zheng et al. based on the observation that the transcription factor *IRF4* is constitutively expressed in Tregs and is necessary for suppression of Th2-mediated activity (56). Similar instances of Tregs’ strategy of co-opting transcriptional pathways characteristic of effector T cells have been observed for Th1 and Th17 suppression (57, 58). Though the exact mechanism remains elusive, it has been posited that Tregs use components of effector T-cell differentiation as a strategy to ‘program’ their deployment to specific inflammatory microenvironments (59). Our data is consistent with this model based on two observations. First, the Th2 signature we identified in Tregs contains high expression of *IKZF2* indicating that this population of Tregs, that is expanded in JDM, originate from the thymus (42–44). Second, the same subset of Tregs upregulate transcriptomic and proteomic expression of CCR4–paralleled by increased expression of CCR4 in CD4^+^ effector T cells– which is preferentially expressed in Tregs recruited to the skin (45). Thus, we speculate that this expanded population of Th2-like Tregs in JDM could represent a peripheral response to site-specific Th2-mediated inflammation in disease-affected tissue potentially driven by autoantibody-producing extra-follicular B-T cell interactions. Future studies should focus on validating expansion of these CCR4^+^ Treg and CCR4^+^ Teff populations in the blood of a larger cohort of JDM patients and functional in vitro studies of Tregs isolated from JDM tissue to determine their suppressive capacity as Treg dysfunction lies at the heart of many autoimmune conditions. Spatial profiling of target tissues would also be insightful to determine the architectural organization, presence of tertiary lymphoid structures, and cell-cell interactions of lymphocytes in JDM muscle and skin.

The observation that type I IFN responses are seen in every immune cell type and increase with clinical metrics of disease activity adds to the growing body of work suggesting that disease activity in JDM correlates with this transcriptional signature (16, 51, 60). However, given the time and cost, it remains infeasible to use transcriptomic sequencing as a lab-based clinical diagnostic tool. Our data points to surface expression of SIGLEC-1 in monocytes is a composite measure of the IFN gene signature in JDM and correlates to clinical disease activity. Together with a recent independent study of JDM in a cohort of similar size, we provide external validation that SIGLEC-1 is a suitable biomarker for disease monitoring to pursue in larger immunophenotyping validation studies given the lower cost and ease of implementing screening by flow cytometry (31). Although profiling blood limits the mechanistic insight compared to skin or muscle, it is a more suitable sample type for biomarker discovery, particularly in a pediatric disease that requires longitudinal monitoring. Importantly, we show that SIGLEC-1 directly reflects the IFN gene signature using paired gene and protein expression measurements, strengthening support for its use as a biomarker. Finally, our work highlights that the IFN gene signature is heterogeneous in JDM. This suggests that SIGLEC-1 levels could also be used to stratify and nominate patients who may benefit from IFN-directed therapies. Further study of this biomarker, and the role of SIGLEC-1 in disease, is an important step toward precision care of JDM.

Finally, we show that integrative approaches like DECIPHERseq can be used to consolidate disparate findings into a systems-level understanding of how interactions among cell states could manifest in disease. Here, our network analysis revealed that a module of hyper-activated IFN response across cell types is coordinated with dysfunction in ribosomal biogenesis, protein processing, and the regulation of cell death that is also shared across many cell types. This model contextualizes recent work that has identified ribosomal dysfunction in NK cells as a disease signature in JDM but also raises the possibility that defective translational machinery is not unique to that cell population (15, 23). Given that Type I IFN directly promotes the activation and proliferation of NK cells (61–63), we speculate that NK cells in JDM are unable to properly translate cytolytic protein machinery required for effector function in response to IFN signaling, potentially perpetuating the IFN response. Similarly, the shared program between CD8T, gdT, and NK cells that describes regulation of cell death and cellular stress response suggests a common dysfunction across cytotoxic cell populations in JDM. Given the importance of cytotoxic cells in clearing cellular debris including autoantigenic neutrophil extracellular traps shown to be pathogenic in JDM (12), dysfunctional cytotoxic populations could result in accumulation of such debris thereby triggering an autoimmune response mediated by lymphocytes. Future work is required to parse the mechanism by which these cells acquire these defects and characterize downstream effects in other immune populations.

These findings should be interpreted in the context of the study’s limitations. First, despite being the largest single-cell study in JDM to date, sample numbers are limited when compared to cohorts used in immunophenotyping studies. Larger studies are necessary to validate SIGLEC-1 expression as a biomarker of IFN-mediated disease activity and confirm expansion of CCR4^+^ CD4^+^ T cell subsets as well as capture the phenotypic heterogeneity of JDM. Additionally, this study lacked data from matched JDM skin and muscle which would have enabled insight into how dysregulated cell states in blood might influence local microenvironments in disease-affected tissue.

In summary, using CITEseq to profile compositional and functional imbalance throughout changes in disease activity, we provide a comprehensive map of the coordinated immune dysregulation underlying JDM. We identify a candidate biomarker that reflects the broadly shared overactive IFN response and provide support for its potential utility in disease management. Furthermore, we reveal novel cell states potentially upstream of this IFN signature that generate new hypotheses for the role of extra-follicular interactions in disease pathogenesis, drawing parallels to other autoimmune diseases. Importantly, these findings pose a new paradigm to how we approach JDM treatment. The dysregulation of processes simultaneously with hyperactivation of other cell states necessitates that we identify therapeutic strategies that restore balance to the dynamic interactions between immune populations rather than simply turning off a set of pathways. Taken together, our work sets the stage for improving clinical management of JDM by providing a foundation for systems-level inquiry into the cellular basis of this disease. More broadly, application of a similar analytical strategy could provide insight into the immunologic basis of other childhood-onset autoimmune diseases characterized by a type I IFN gene signature.

## Methods

### Study Cohort & Sample Processing

Patients were recruited to the Juvenile Myositis Precision Medicine Biorepository between 2018 and 2021 and underwent informed consent. This study was approved by the UCSF IRB. Clinical data was collected by study investigators and recorded in a secure REDCap database. Treatment-naive JDM was defined as a new diagnosis of JDM as deemed by the treating clinician with no systemic immune suppressant use in the prior 4 weeks. Inactive JDM was defined as normal CK, MMT8≥78 and Physician Global VAS score<0.5 to reflect PRINTO clinically inactive disease definitions but with some modifications based on the data available. Active disease was defined as Physician Global VAS score≥0.5 and taking immune suppressive medication. Healthy controls were enrolled who had no prior autoimmunity, no known or suspected genetic disorders, immunodeficiency, active cancer, or history of organ or bone marrow transplantation, no infection or antibiotics in the prior 4 weeks, no chronic systemic immunomodulatory medication use and no vaccinations in the prior 6 weeks. Peripheral blood samples were collected at each study visit and processed by the Pediatric Clinical Research Core Sample Processing Lab. PBMCs were collected in SepMate tubes (n=9) using Ficoll separation or CPT tubes (n=15), isolated per manufacturer’s guidelines, and cryopreserved in liquid nitrogen.

### CITE-seq of human PBMCs

Our experimental protocol followed protocol from our previous study (16) with certain modifications to account for confounding time-related and batch effects. Note these experiments were carried out using early access kits from BD Genomics before the implementation of commercially-available single-cell protein/RNA assays (e.g. Feature Barcoding, 10x Genomics; BD Abseq, BD Genomics, Supplemental Table 4), and researchers are recommended to use those newer solutions for any follow-up studies as the techniques and reagents have been refined. PBMCs from 27 distinct samples were gently thawed in a 37°C water bath and re-suspended using a pipette set to 1 mL. Cell counts and viability were determined using a Cellometer Vision (Nexcelcom) with AOPI staining (Nexcelcom cat. CS2-0106-5ML). Cells were multiplexed into four pools: one “cross pool” with all samples that consisted of only one time point and three pools consisting of longitudinal samples. Longitudinal samples from the same individual were assigned to separate pools to enable genetic demultiplexing. After pooling, cells were resuspended in 90 μl of 1% BSA in PBS and Fc blocked with 10 μl Human Trustain FcX (Biolegend cat. 422302) for 10 minutes on ice then stained on ice for 45 minutes with a pool of 268 antibodies in 100 μl, for a final staining volume of 200 μl. Antibodies were pooled on ice with 2.2 μl per antibody per 1×106 cells (BD Genomics). Cells were quenched with 2 ml 1% BSA in PBS and spun at 350xg for 5 minutes and further washed two more times with 2 ml of 1% BSA in PBS. After the final wash, cells were resuspended in 100 ul and strained through a 40 μM filter (SP Bel-Art cat. H13680-0040). Each longitudinal pool was split across two 10X lanes while the “cross pool” was split across six 10X lanes (6 wells total, 5×10^5^ cells/well). The 10x Chromium was run and post-GEM RT and cleanup were done according to manufacturer’s protocol (10X Genomics 3’ Kit V3). Starting at cDNA amplification, modifications to the protocol were made: 1 μl of 2 μM additive primer (BD Genomics, beta kit) specific to the antibodies tags was added to the amplification mixture. During the 0.6X SPRIselect (Beckman Coulter, B23318) isolation of the post-cDNA amplification reaction cleanup, the supernatant fraction was retained for ADT library generation. Subsequent library preparation of the cDNA SPRI-select pellet was done exactly according to protocol, using unique SI PCR Primers (10X Genomics). For the ADT supernatant fraction, a 1.8X SPRI was done to isolate ADTs from other non-specifically amplified sequences, followed by sample index PCR. Sample index PCR for the ADTs was done using the cycling conditions as outlined in the standard protocol (15 cycles) but using unique SI-PCR Primers such that all libraries could be mixed and sequenced together. Subsequent SPRI selection was performed, and all libraries were quantified and analyzed via Qubit 2.0 (Fisher) and Bioanalyzer (Agilent), respectively, for quality control. We sequenced the libraries on 2 lanes of a NovaSeq S4 (Illumina), aligned using CellRanger (10X Genomics) to generate feature barcode matrices.

### Sequencing data pre-processing and integration

Data was demultiplexed using genotypes with demuxlet (64) and doublets were filtered using DoubletFinder (65). Next, the data were filtered to remove genes with < 3 cells. Additional filters were applied to the cells, removing cells with greater than 5000 ADT counts to avoid antibody aggregates and with >60% ribosomal or >15% mitochondrial DNA (mtDNA) reads. For the ADT data, cells were additionally filtered to remove those with fewer than 70 antibodies detected, and with any antibody isotype control measurements greater than 50. To remove background ambient RNA signal, we ran SoupX separately on each of the six RNA libraries and then merged them (66). Aggregated data was log-normalized and scaled, regressing out percent mtDNA, percent ribosomal DNA, and cell cycle (S, G2M) (67). Data was then integrated with Harmony, with 20 max iterations and 30 max iterations per cluster (68).

DSB was run on all six ADT libraries individually, using default parameters except for more stringent quantile clipping (0.01, 0.99) (69). The background distribution of empty droplets was defined as suggested in the DSB vignette. Isotype controls were then removed from the dataset, and RPCA was used to integrate the DSB-normalized ADT data across libraries. Following RPCA, the data was re-scaled and cell cycle scores (S, G2M genes) and the number of ADT counts and features were regressed out. The harmonized RNA and RPCA corrected ADT were combined using Weighted Nearest Neighbors, with default parameters except for prune.SNN = 1/20. Leiden clustering was run on the resulting graph (method = igraph), at a 1.4 resolution (70). Two clusters were removed with low to no expression of ADT and the object was reclustered with the same parameters. The Seurat function ‘FindAllMarkers’ was used to identify the top 5 markers per cluster.

We removed an additional 3 clusters: 2 were very small clusters with a transcriptomic profile consistent with doublets (original Leiden clusters 26 and 29, Supplementary Figure 16), and 1 diffusely expressed cluster (original Leiden cluster 19). We further sub-clustered 3 clusters that expressed genes representative of more than one cell type: original Leiden clusters 16, 17 and 23. Sub-clustering was performed using Seurat’s ‘FindSubCluster’ function using the lowest possible resolution to divide the population into two clusters. Based on minimal transcriptional differences between them, original Leiden clusters 1, 5, 9, 11 and 15 were merged into a single CD4^+^T naïve population, clusters 3 and 10 into a single naive CD8^+^T population, and cluster 7 and part of the subsetted cluster 23 CD56^dim^ NK population. Due to interpersonal heterogeneity in monocytes, all CD14^+^ monocyte clusters were merged into one CD14^+^ monocyte population (71, 72).

While annotating, we discovered that the FOXP3-signature normally attributed to Tregs was only present in a subset of the cluster and FindSubCluster did not appropriately isolate the *FOXP3^+^*cells. We therefore subsetted the cluster and re-ran ‘FindVariableFeatures’, ‘ScaleData’, ‘RunPCA’, ‘FindNeighbours’, ‘FindClusters’ with the Louvain algorithm and a resolution of 0.8, and ‘RunUMAP’. This enabled us to subset a smaller group of cells with a statistically significant expression of *FOXP3* compared to other clusters using FindAllMarkers, which we hence annotated T regulatory cells. Annotation of the remaining clusters was performed using both canonical gene and protein markers. One B cell population consisted almost solely of cells from two donors. This was annotated as B_naive4, and was not used in downstream analysis, but included in UMAPs.

### Cell type proportion analysis

Cell type proportion was calculated as the proportion of each cell type for each individual and was compared for: treatment-naive JDM compared to HC, treatment-naive JDM compared to inactive JDM and inactive JDM compared to HC using Kruskal-Wallis test with Dunn’s post-test. To determine the association between cell abundance and disease activity, the Spearman correlation coefficient between cell type proportion and physician global VAS scores was calculate and p values were adjusted using BH.

### Differential gene and protein expression analysis

The DGE and DPE analysis were completed using DESeq2. Size factors were set using the function ‘computeSumFactors’ from the scran package. We used the default settings for single cell data, namely test=‘LRT’, useT = T, minmu = 1e-6, fitType = ‘glmGamPoi’, and minReplicatesForReplace = Inf in the ‘DESeq’ function. Batch was included as a co-variate using the ‘reduced’ argument. We filtered genes and proteins that were not expressed in at least 5% of cells and analyzed only cell types where there were at least 100 cells in each group. We used cutoffs of |LFC| ≥ 1 for genes, |LFC| ≥ 0.5 for proteins, and BH p < 0.05. Over-representation analysis was performed on up- and downregulated genes per cell type using the clusterProfiler package with GOBP as reference and adjusted p < 0.05 (). For the PD1/CD45R0-subanalysis, we compared groups using Seurat’s FindMarkers with test.use = ‘MAST’, latent.vars = ‘well’, |LFC| ≥ 0.5, and BH p<0.05.

### Identification of global IFN signature

We created a list of IFN genes by excluding cell types with less than 100 cells in either HC or treatment-naive JDM and then collected genes differentially expressed in at least 2 cell types. The average gene expression was calculated using Seurat’s ‘AverageExpression’ function. Expression was averaged per sample for each cell type. The expression was visualized using dittoSeq’s ‘dittoHeatmap’ (73) with default, unsupervised clustering settings of both rows and columns, and the dendrograms ordered using the dendsort package (74). The clustering organized the genes into 7 distinct modules, where Module 1 consisted exclusively of IFN-related genes. Average Module 1 scores for each cell type were then calculated using Seurat’s ‘AddModuleScore’ with default settings. Correlations between disease activity and IFN score was calculated using Spearman correlation and visualized using ggplot2 (75).

### Network inference from RNA data using DECIPHERseq

We applied NMF to the raw RNA count data as implemented in the DECIPHERseq method with default parameters (25). The main output of NMF is a set of two orthogonal vectors: gene loadings that represent how much a given gene contributes to that activity program, and cell loadings that represent how strongly that program is expressed in a given cell. The NMF rank, k, was chosen using the weighted subtrees metric based on phylogenetic clustering, as described by Murrow et al. The final choices of rank k for each cell type were k_B_=17, k_CD4T_=17, k_CD8T_=14, k_gdT_=13, k_Myeloid_=17, k_NK_=11 according to the saturation point in the elbow plots (Supplemental Figure 17). Network clustering was performed on the per-sample averaged program scores with default parameters as described by Murrow et al. The corresponding gene loading vectors for each GEP were analyzed as described by Kotliar et al. to quantify the strength of an individual gene’s contribution to that program, referred to as ‘marker gene scores’ (36). GSEA was performed on the resultant ranked gene lists using the fgsea (76) package in R with GO and Hallmark gene sets. Module themes were assigned by calculating module enrichment p-values using the ‘Get_enrichment_pvals’ function in DECIPHERseq with default parameters. Module and gene set enrichment results were visualized using ClusterProfiler (77).

### Statistics

All statistical analyses and visualization of results were performed using open-sourced R (version 4.2.3). Pairwise comparisons of cell proportions between patient groups were performed using a Kruskal-Wallis test with post-hoc Dunn comparison, with p-values adjusted for multiple comparisons by Holm correction. Significance of Pearson correlations between GEPs used for network construction was calculated using bootstrapping as implemented in DECIPHERseq. Analyses of disease association with GEPs was performed using two-tailed unpaired t-test or ANOVA with post-hoc Tukey test. False discovery rates for GSEA annotation and module enrichment across programs were calculated and corrected at the cutoff FDR < 0.01 as described by Murrow et al. Gene loading similarity was calculated as the Pearson correlation between gene loadings for each activity program and all other activity programs in the same module with p-values calculated by permutation testing. Correlation methods used in specific figures are described in the corresponding legends and in Methods, and significance for statistical tests was set at the threshold P < 0.05.

### Study approval

This study was approved by the UCSF IRB #17-24003. Written informed consent to participate in this study was provided by the participant or the participants’ legal guardian depending on the age of the participant. Assent was obtained when appropriate.

### Data availability

The datasets presented in this study are deposited in the CZ CELLxGENE Discover resource as ‘CITEseq of JDM PBMCs’ (https://cellxgene.cziscience.com/collections/c672834e-c3e3-49cb-81a5-4c844be4a975). The code used for this analysis will be made publicly available on Github at “grabadam-cal/jdm_crosslong” upon manuscript acceptance.

## Author Contributions

GR and CW were assigned co-first authorship because their contributions were deemed equally essential to the study, with GR listed first because she led the writing of the manuscript. JN and SK recruited the patients for the study. GCH, YS, and JN performed the experiments. GCH performed the alignment and sample demultiplexing. EF, GR, and CW performed the integration and quality control. CW performed the annotations and the cell proportion and differential analysis. GR conceptualized and wrote the pipeline for the network analyses; EF ran the pipeline. ZG, MS, and JN provided essential feedback on the analysis strategy and statistical methods. GR, CW, and JN wrote the manuscript; MS and ZG revised the manuscript. JN, MS, and SK conceptualized the study. JN led the project administration and acquired funding for the study. All authors reviewed and edited the manuscript.

## Acknowledgements

The authors would first like to express gratitude for the patients with JDM and their families who contributed their samples and data. We would also like to acknowledge the Institute of Human Genetics Genomics Core at UCSF, Center for Advanced Technology at UCSF, and the UC Berkeley Vincent J. Coates Genomics Sequencing Laboratory. We thank the members of the UCSF Pediatric Rheumatology Division for helping to recruit patients: Emily von Scheven, William Bernal, Michael Waterfield, Erica Lawson, Alice Chan, Geraldina Lionetti, Nicole Ling, Tara Valcarcel, Andy Nguyen, William Soulsby, Julia Shalen, Sara Haro, Tyalor Laflam, and our research coordinators, Bhupinder Nahal, Kathy Nguyen, Katrina Gonzales, Bethany Bautista & Zian Zheng. JN was supported by the Cure JM Foundation and the Doris Duke Physician Scientist Fellowship, Grant 2019124. GR was supported by the NIH ipCBS Fellowship, NIH T32-GM008155, the Rheumatology Research Foundation and in part by grants from the NIH (R01GM135462 and R33CA247744) and the UCSF Center for Cellular Construction (DBI-1548297), an NSF Science and Technology Center. CW was supported by the DARE Fellowship, Fabrikant Aage Lichtingers Foundation, Director Ib Henriksens award, Knud Hoejgaards Foundation, Jorcks Foundation, and the Denmark-America Foundation. GCH was supported by the NSF Graduate Research Fellowship. MS, CJY, and EF were supported by P30-AR070155. Z.J.G. is a Chan Zuckerberg BioHub San Francisco Investigator.

