## Supplemental Figures for "Coordinated immune dysregulation in Juvenile Dermatomyositis revealed by single-cell genomics"

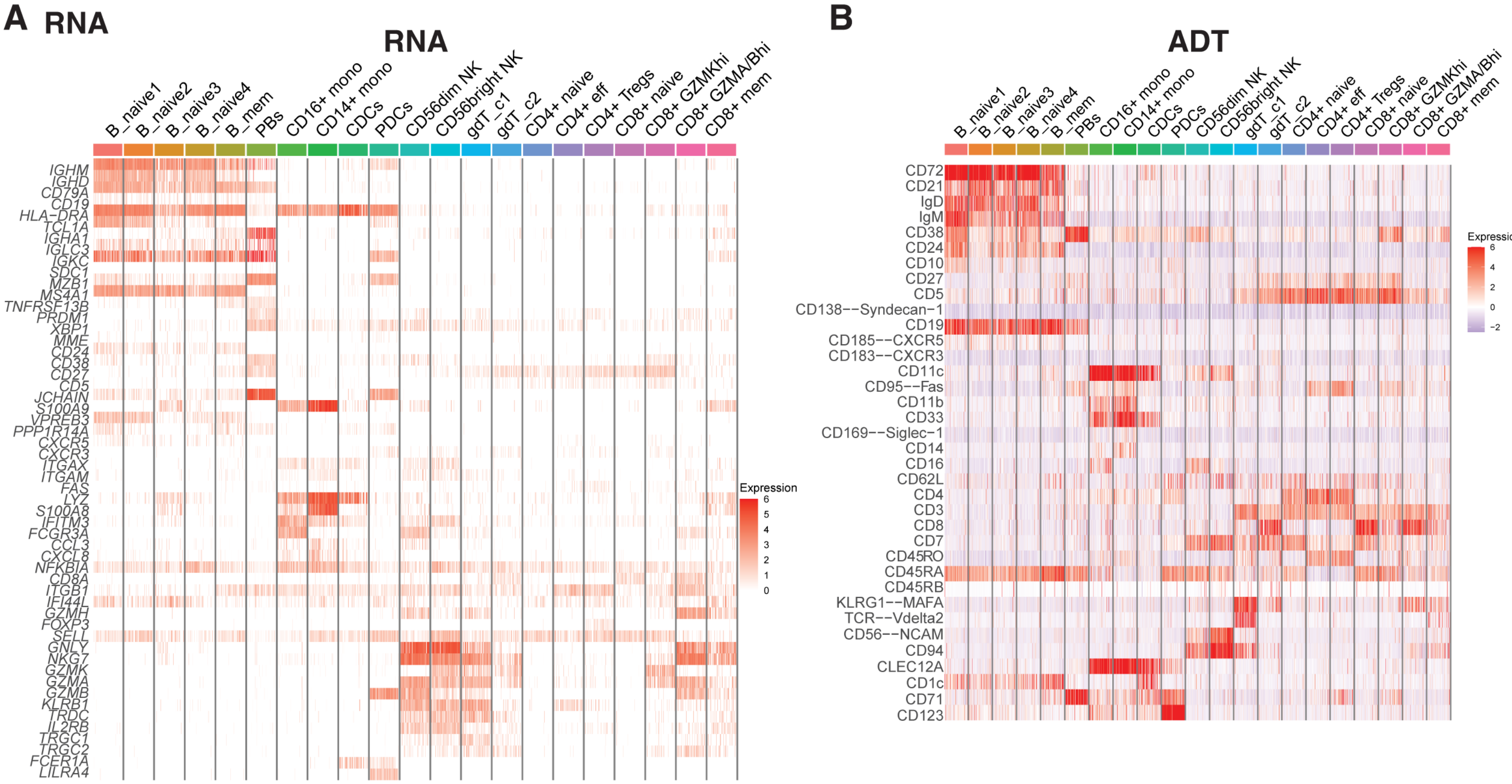


**Supplemental Figure 1.** Canonical RNA **(A)** and surface protein **(B)** markers for clusters in wnnUMAP shown in Figure 1.

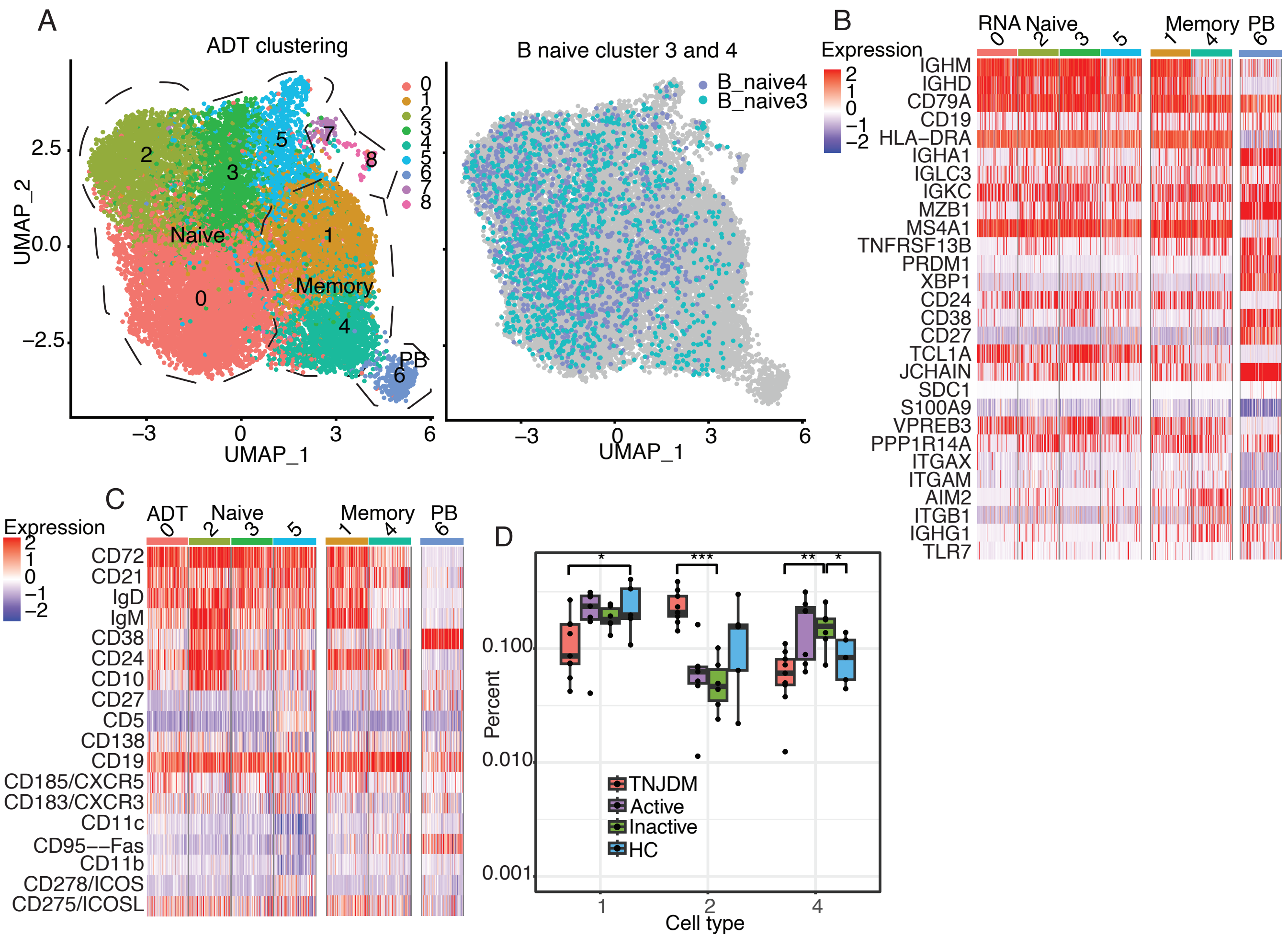


Supplemental Figure 2. Analysis of reclustered B cells based on ADT measurements alone.This re-clustering was completed on the DSB-corrected ADT assay using the Seurat workflow. FindClusters was run using a resolution of 0.4 and with the Leiden algorithm. FindUMAP was run using default settings. (A) UMAP of subsetted B cells where Clusters 0, 3 and 5 corresponded to naïve B cells, cluster 2 to an immature naïve B cell population akin to “B_naive1” in the first analysis, and clusters 1 and 4 to IgM+IgD+ memory and IgM-IgD- memory B cells, respectively, defined by TNFRSF13B (TACI) expression. Cluster 6 consisted of plasmablasts. Clusters 7 and 8 contained few cells with a high expression of platelet and red blood cells specific genes, and were excluded from further analysis. B_naive3 and B_naive4, clusters driven by RNA signatures, do not form specific clusters and patient-specific clustering is resolved. Canonical RNA (B) and ADT (C) markers for reclustered B cells. (D) Compositional analysis using these ADT clusters verified an increase in the proportion of immature naïve B cells (IgM+IgD+CD24+CD38+CD10+) from cluster 2 in TNJDM as compared to HC. This analysis also found a significant decrease in proportion of memory B clusters 1 and 4, in TNJDM as well as an increase in the proportion of IgM-IgD- memory B cells (cluster 4) in inactive JDM.

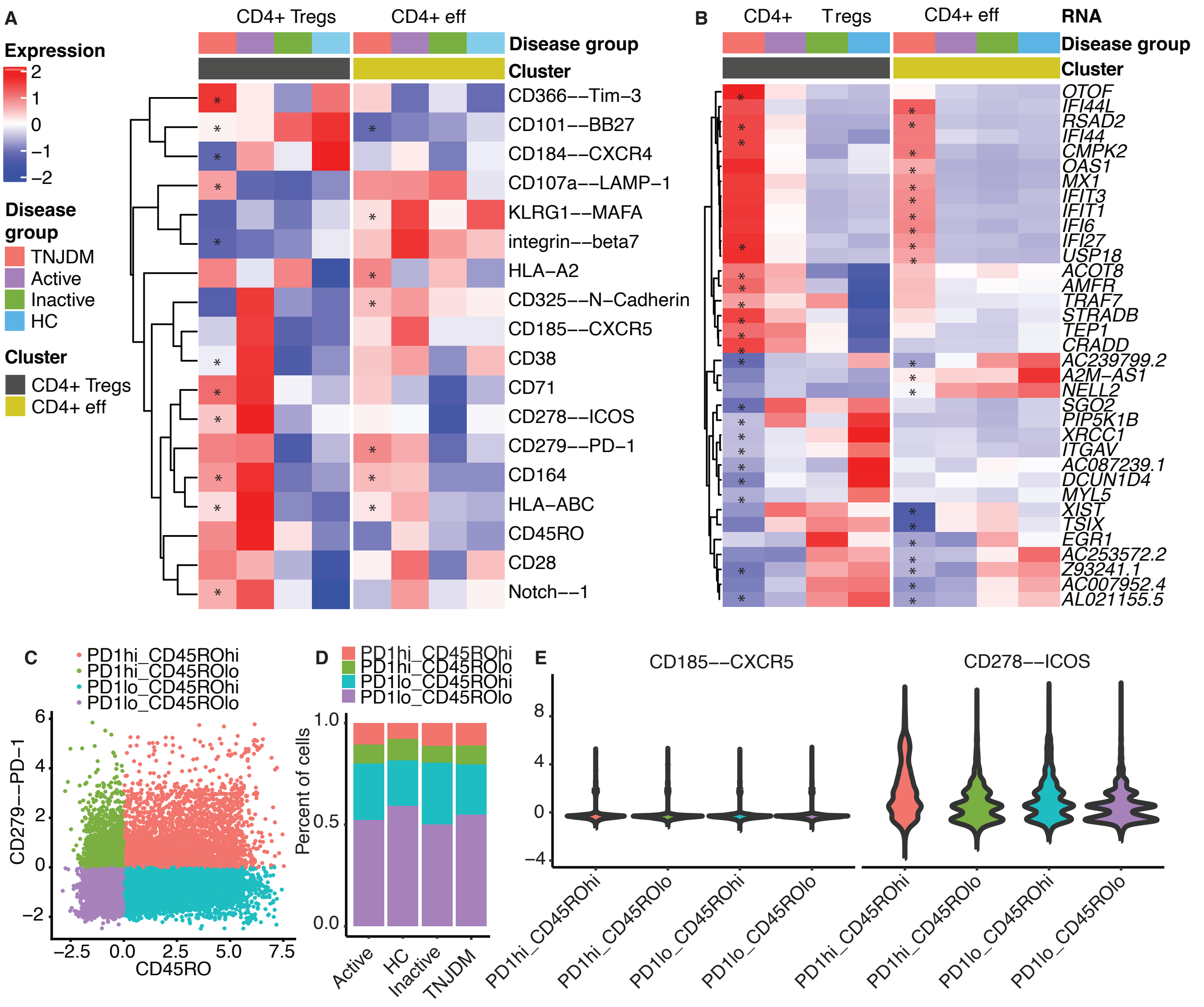


**Supplemental Figure 3. (A)** Heatmap of select protein ADTs. Asterisks mark signifcant comparisons between TNJDM and HC for Tregs and CD4+ effector T cells with an absolute logfold change ≥0.5 and an adjusted p-value < 0.05. **(B)** Heatmap of top 10 and bottom 10 differentially expressed genes between TNJDM and HC for Tregs and CD4+ effector T cells with an adjusted p-value ≥ 0.05. Asterisks mark significant comparisons. **(C)** Dot plot of CD4+ T cells showing expression of CD45R0 and PD-1. **(D)** Bar plot showing percentages of PD1/CD45R0-expression groups per disease group. **(E)** Violin plots showing expression of CXCR5 and ICOS per PD1/CD45R0-expression rgoup

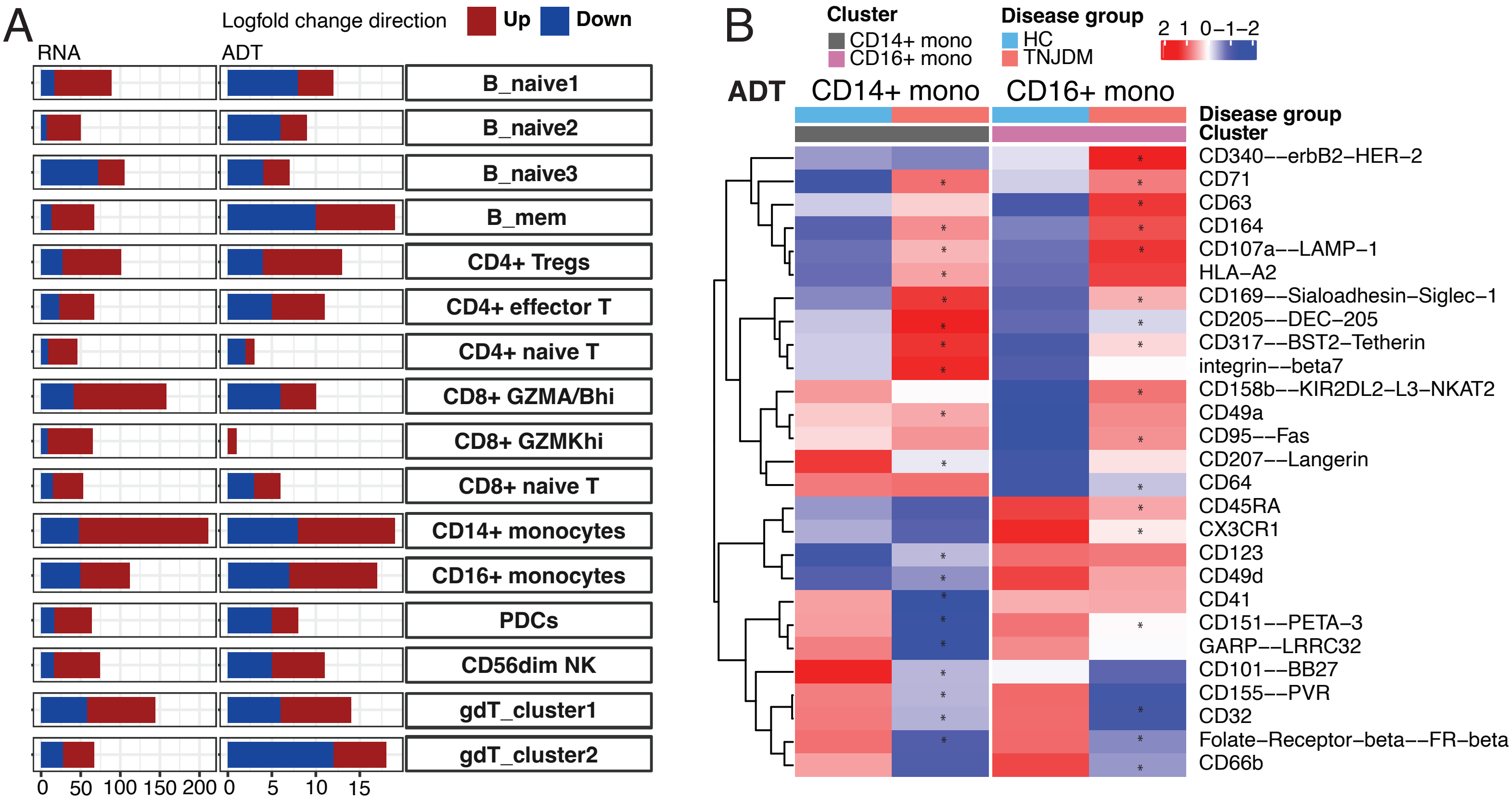


**Supplemental Figure 4. (A)** Differential analysis between treatment-naive JDM and HCs for each cell type. **(B)** Differential analysis of surface proteins (TNJDM vs. HC) for monocytes. Asterisks indicate significant differential expression (BH adjusted $p < 0.05$).

**A**

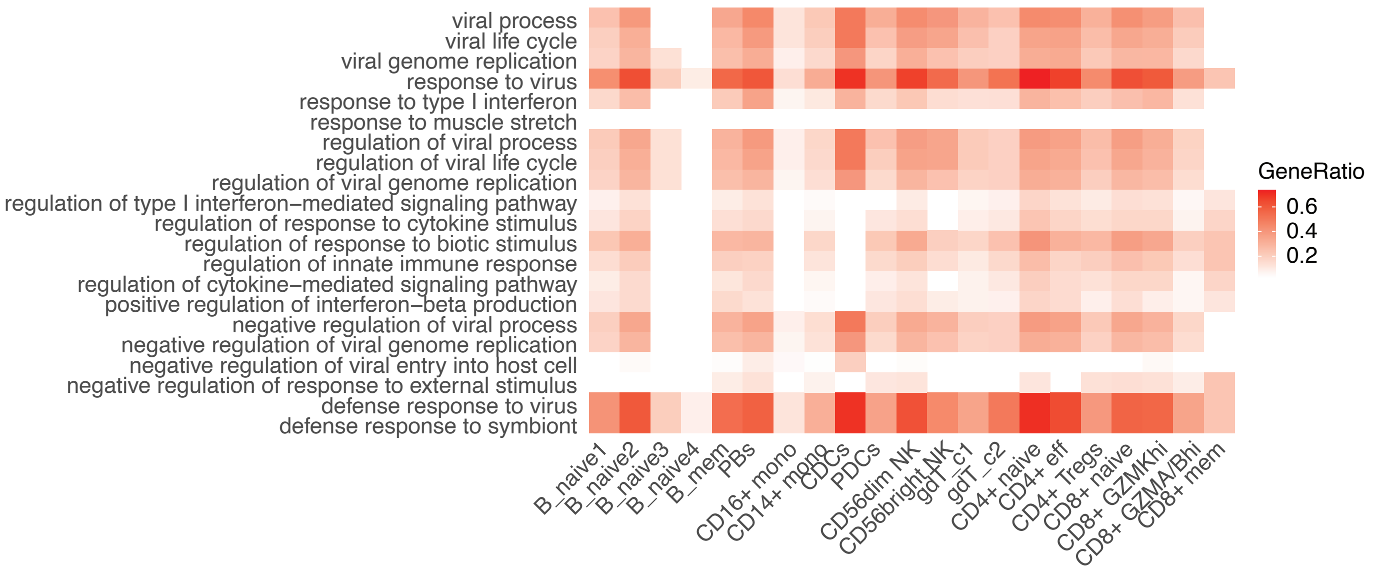


**Supplemental Figure 5.** Heatmap of enriched GO terms related to Type I Interferon Response from GSEA with FDR<0.01.

A

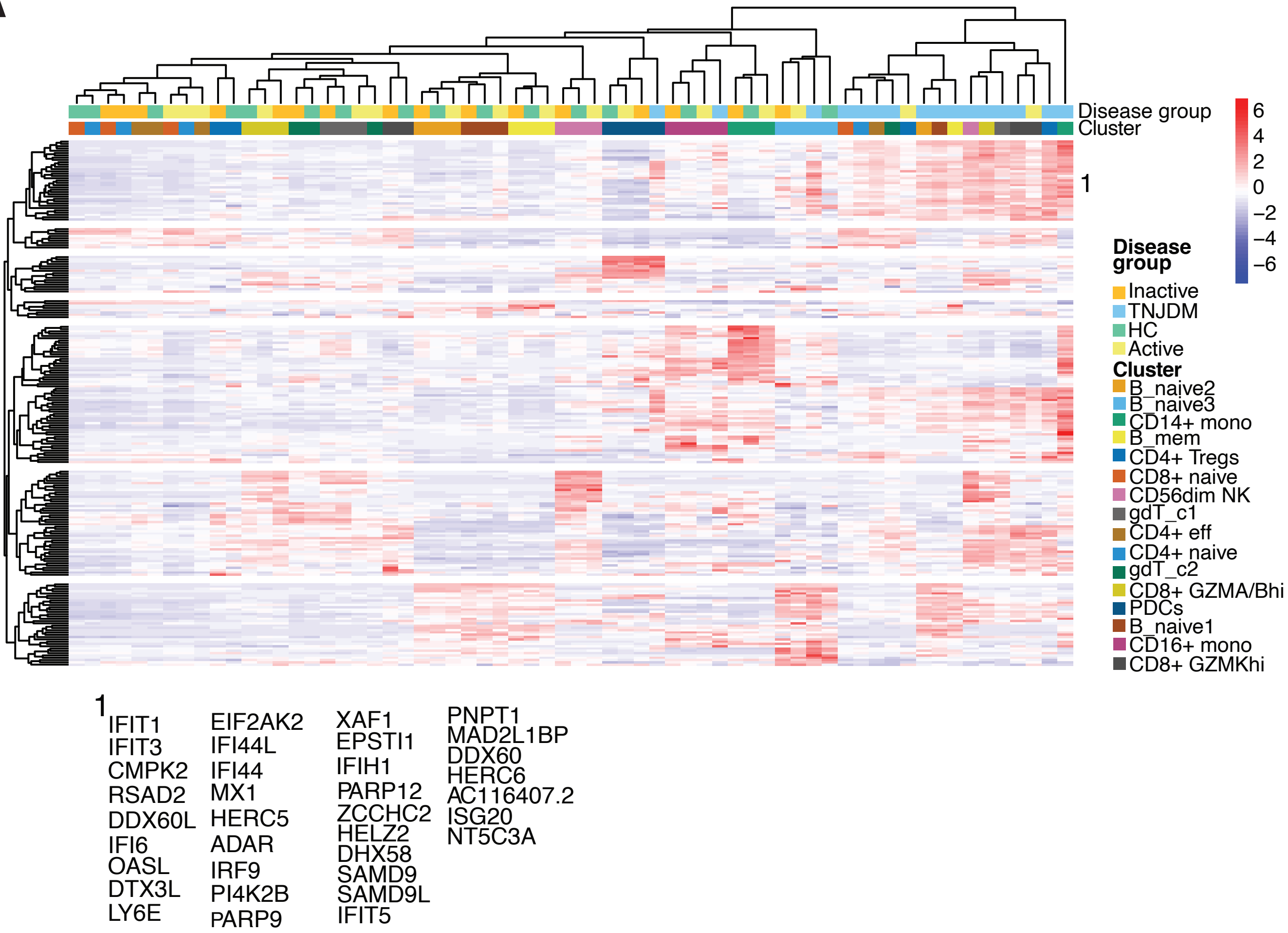


**Supplemental Figure 6. (A)** Heatmap of differentially expressed genes between TNJDM and HC from all cell types clustered by expression likeliness. The genes from cluster 1 were used to calculate the IFN score

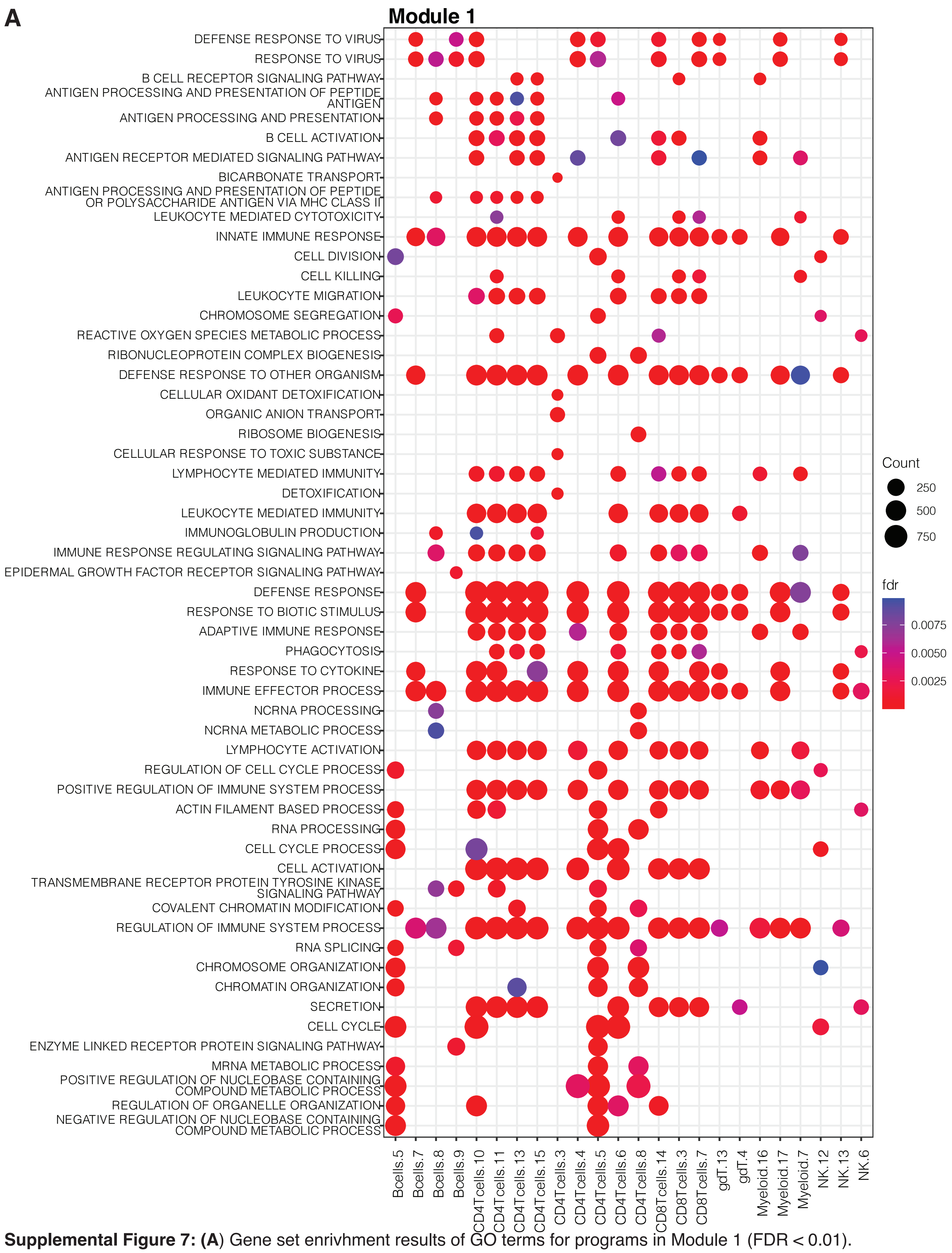


**Supplemental Figure 7: (A**) Gene set enrivhment results of GO terms for programs in Module 1 (FDR < 0.01).

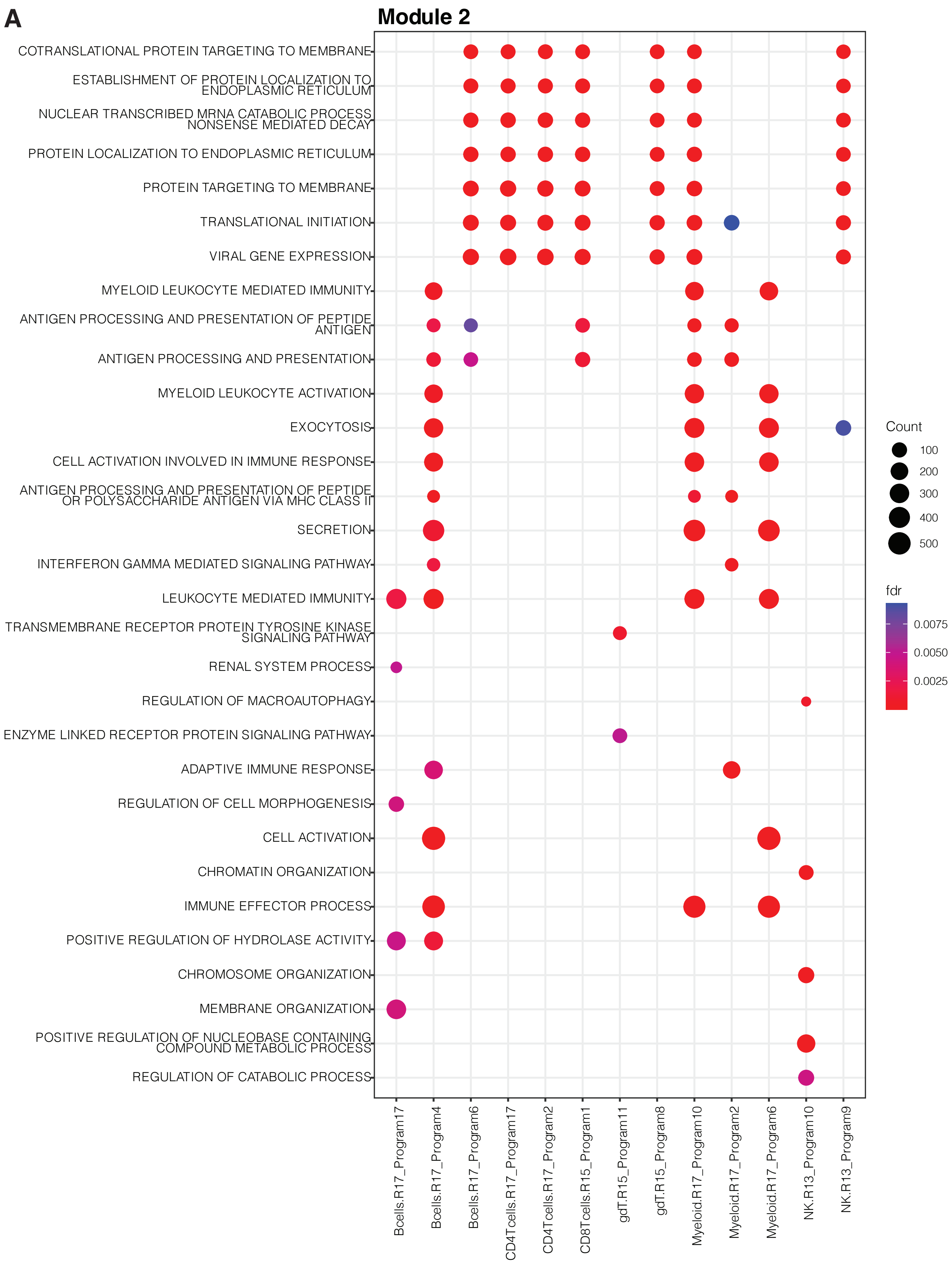


**Supplemental Figure 8: (A)** Gene set enrivhment results of GO terms for programs in Module 2 (FDR < 0.01).

A

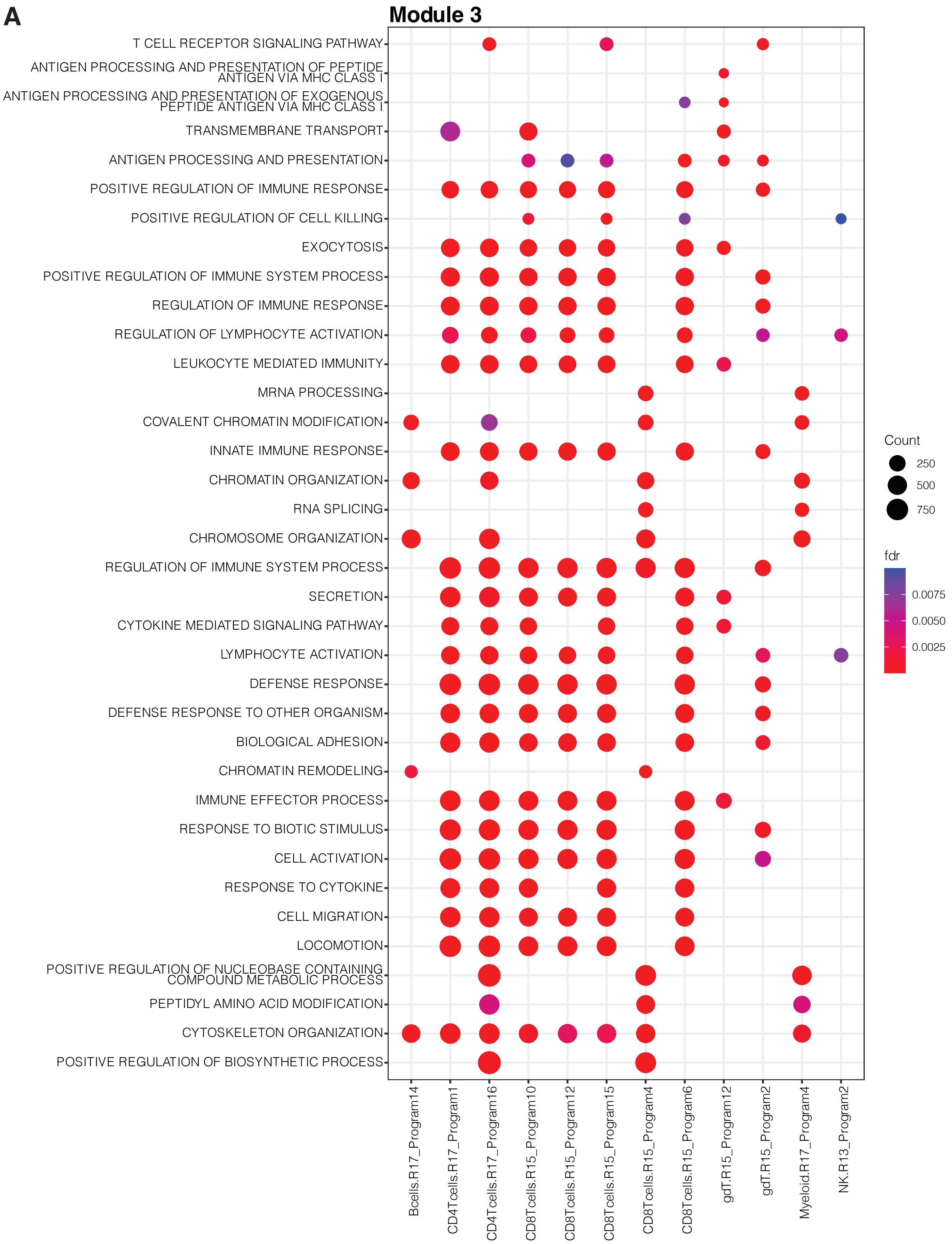


**Supplemental Figure 9: (A)** Gene set enrivhment results of GO terms for programs in Module 3 (FDR < 0.01).

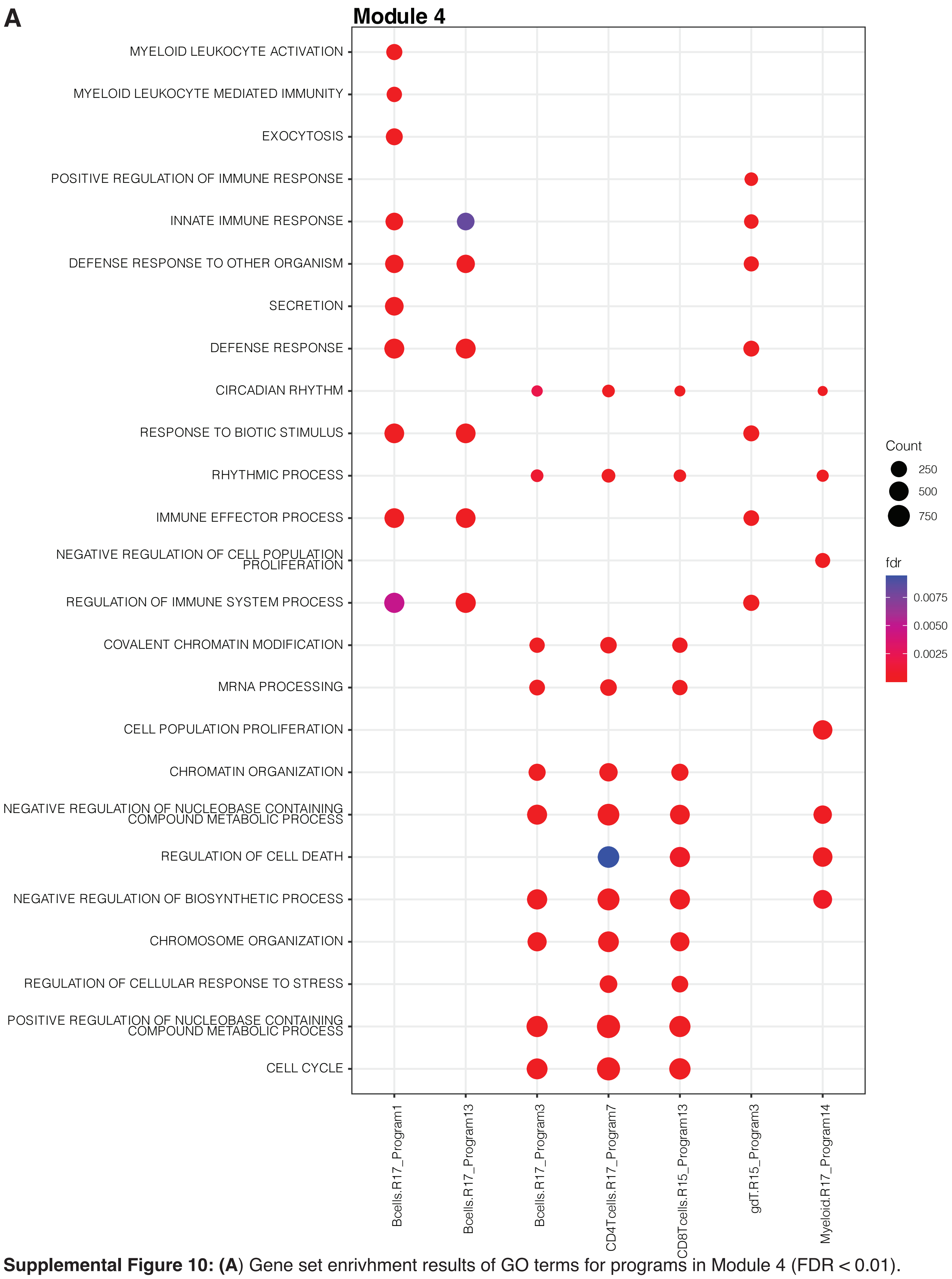


**Supplemental Figure 10:** (**A**) Gene set enrivhment results of GO terms for programs in Module 4 (FDR < 0.01).

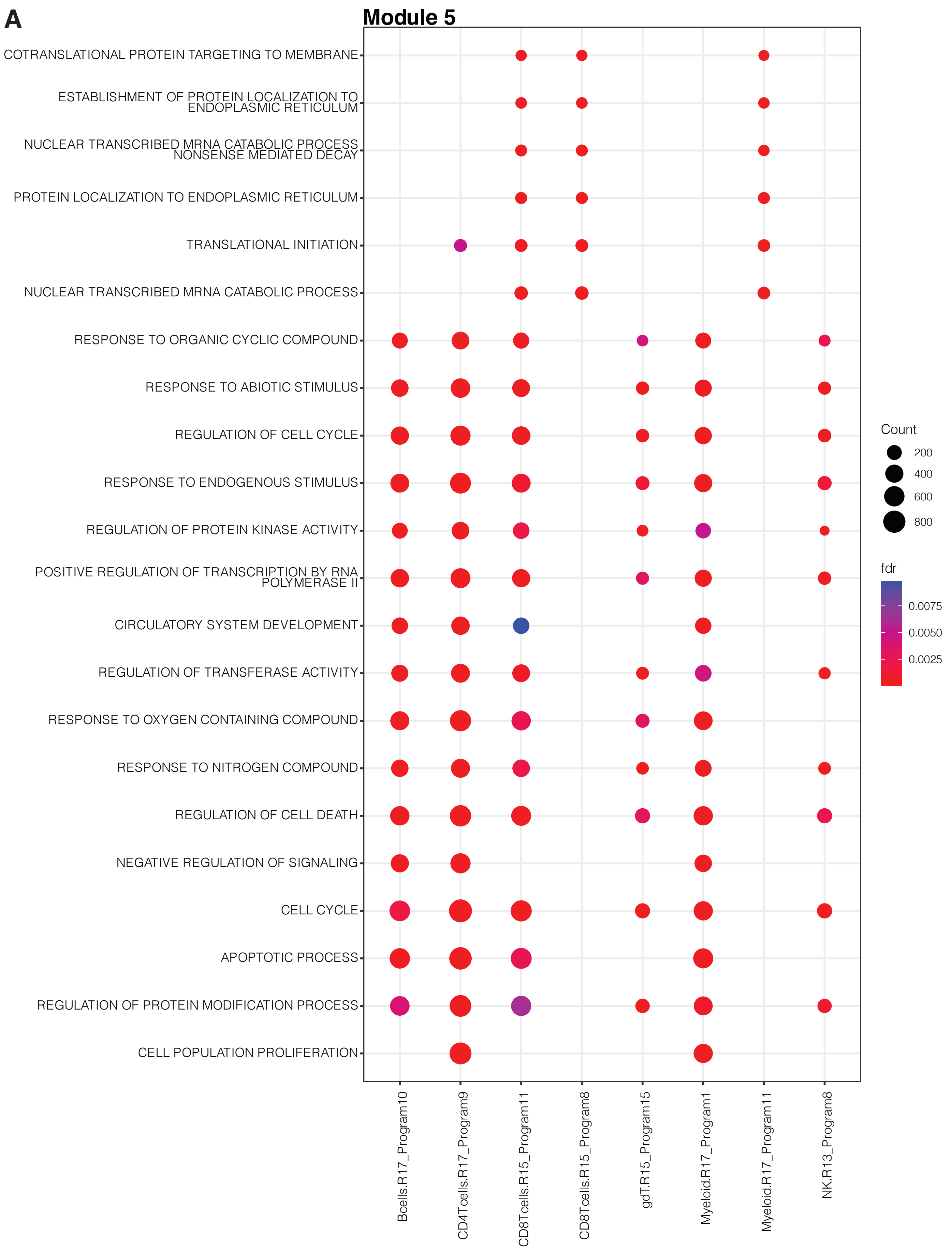


**Supplemental Figure 11: (A)** Gene set enrivhment results of GO terms for programs in Module 5 (FDR < 0.01).

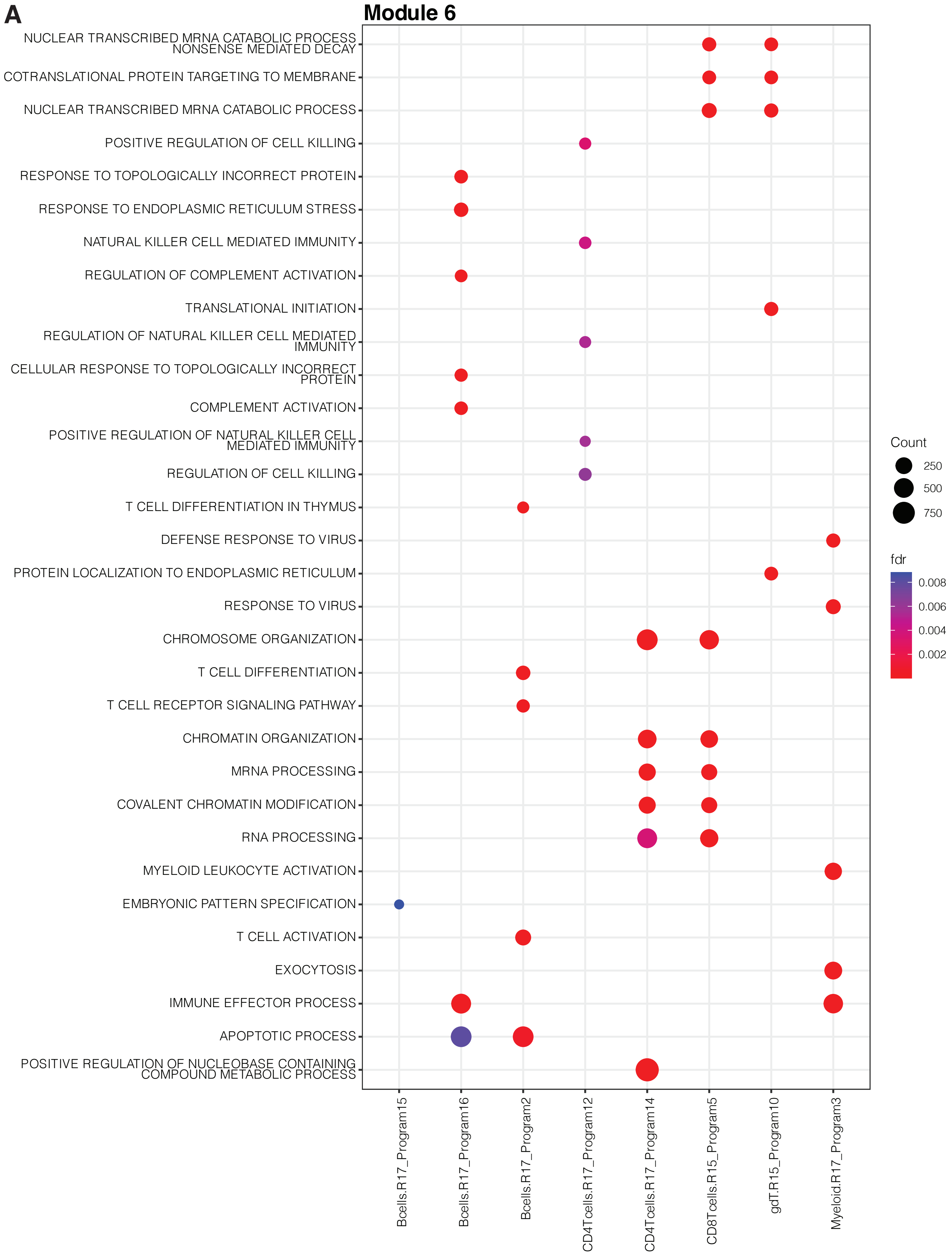


**Supplemental Figure 12: (A)** Gene set enrivhment results of GO terms for programs in Module 6 (FDR < 0.01).

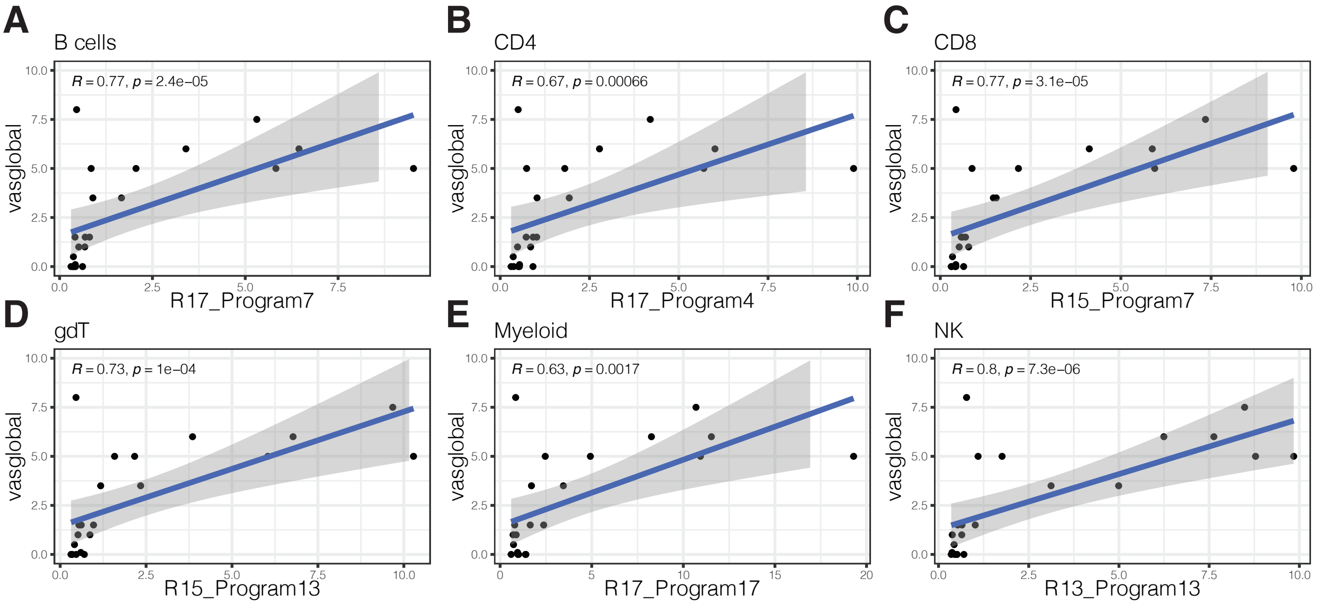


**Supplementary Figure 13: (A-F)** Scatter plots showing mean sample expression (n=27) of type I interferon response programs in each corresponding cell type (Pearson).

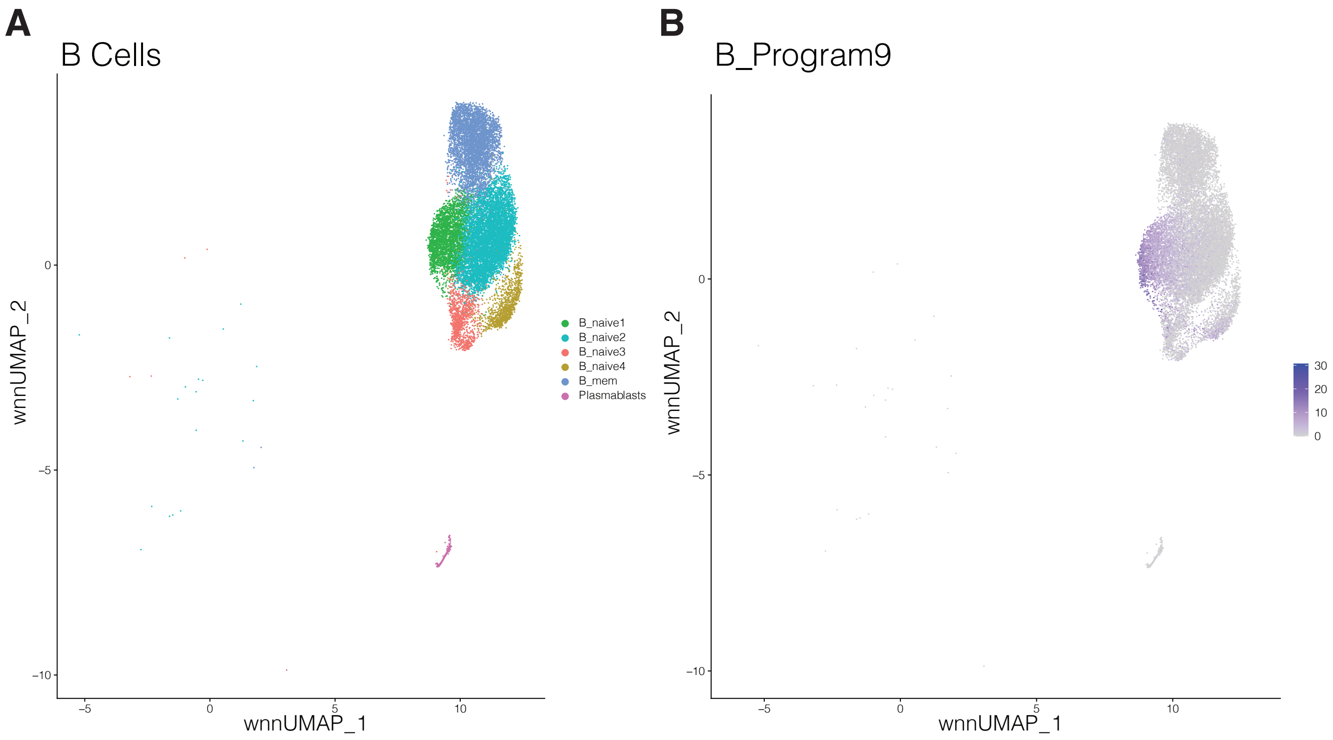


**Supplemental Figure 14 (A)** wnnUMAP of B cell subsets **(B)** wnnUMAP showing expression of program B9 in B cells.

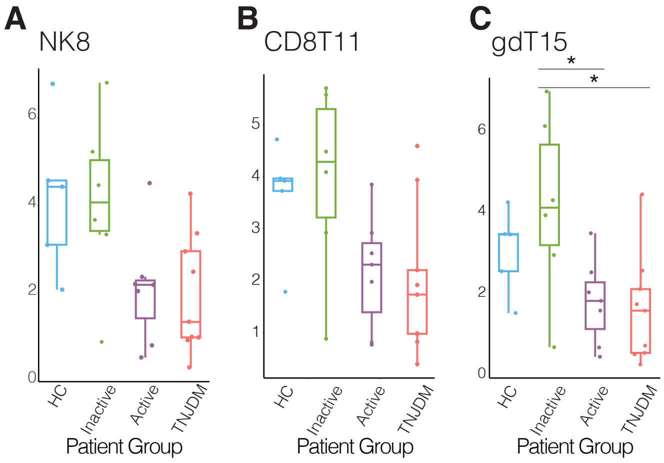


**Supplemental Figure 15 (A-C)** Mean patient expression of disease activity associated programs (4-way ANOVA, $p<0.05$) in Module 5 (*$p<0.05$ Post-hoc pairwise Tukey test).

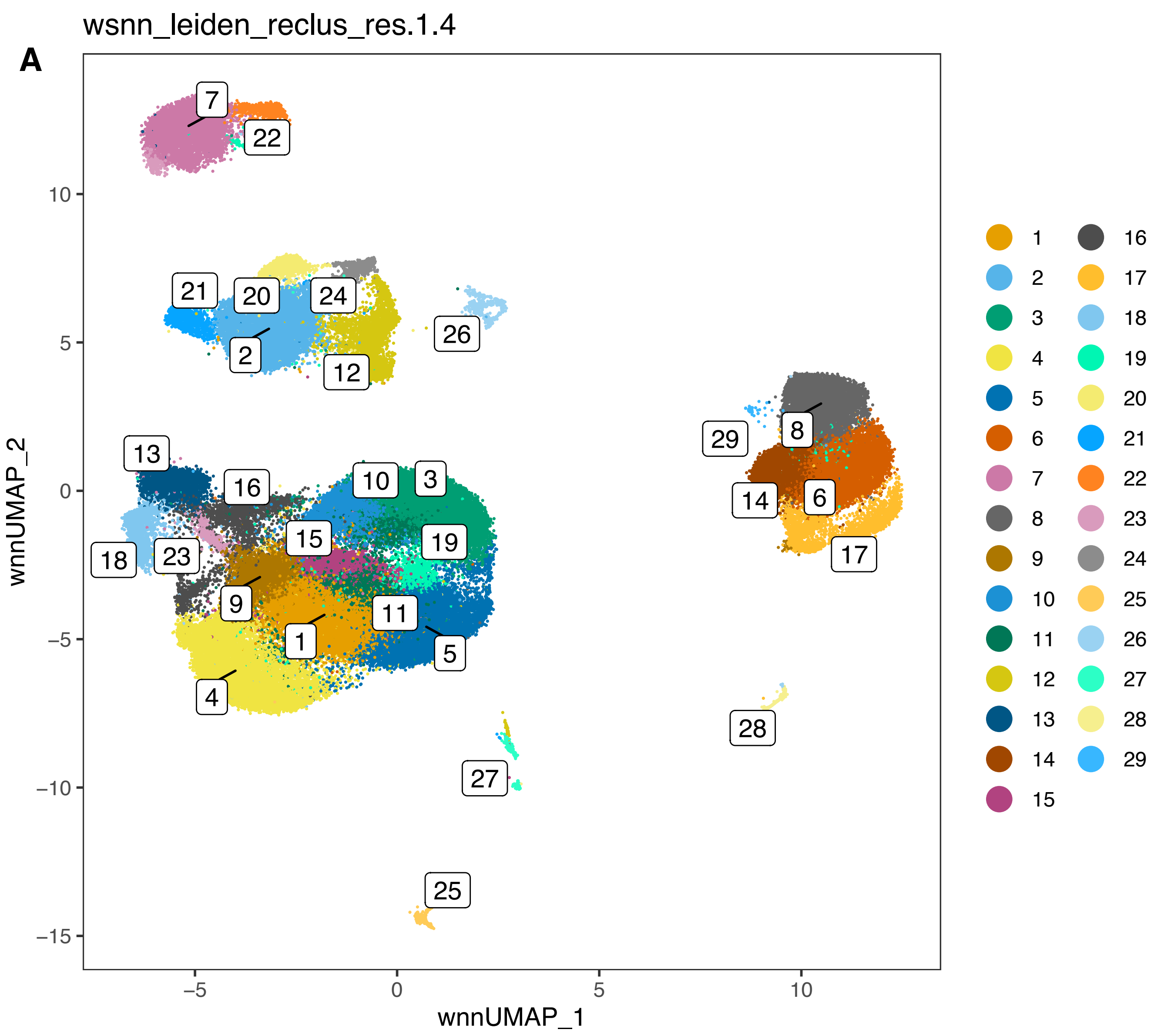


**Supplementary figure 16. (A)** Original wnnUMAP, using Leiden clustering with a resolution of 1.4

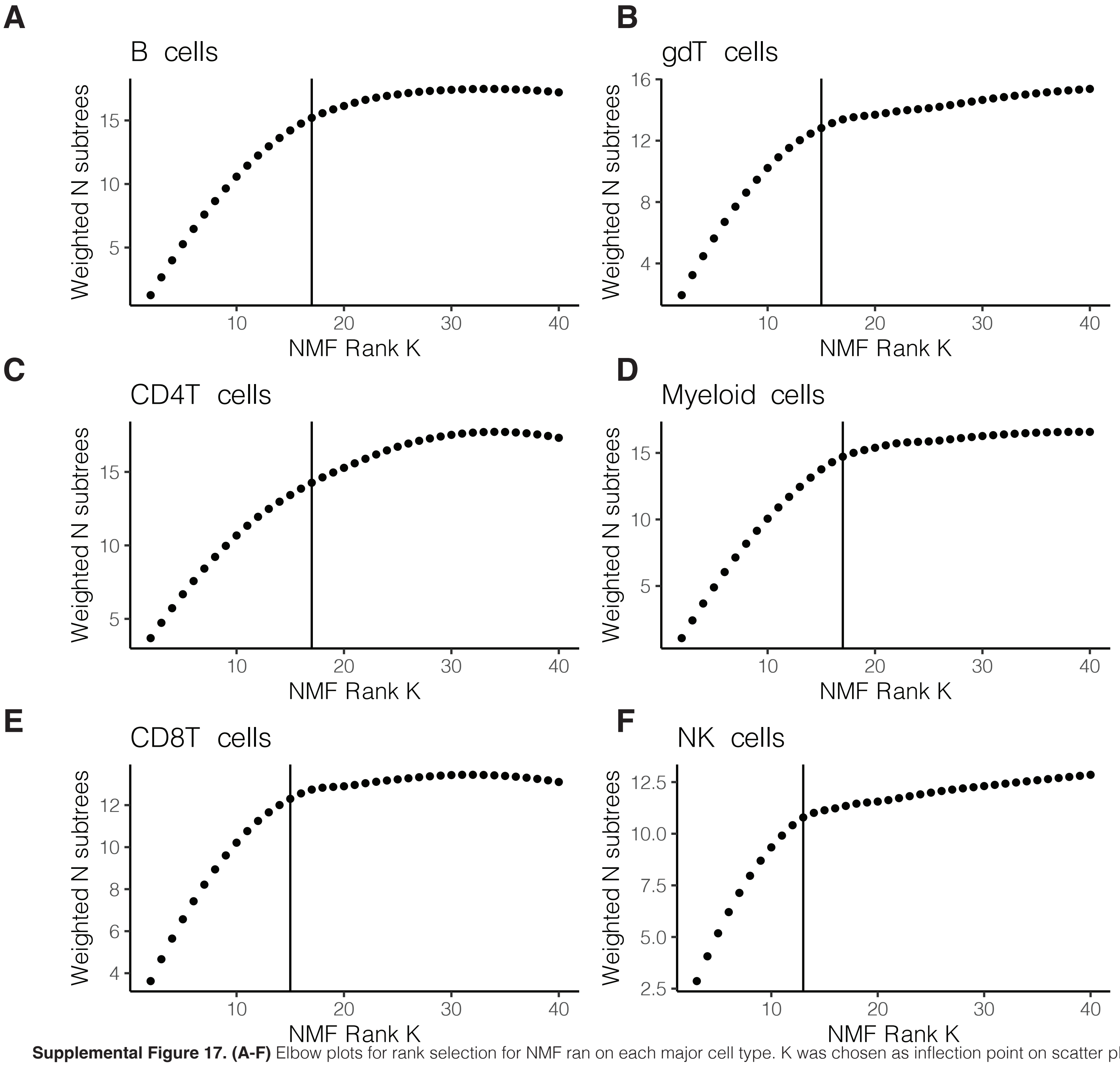


**Supplemental Figure 17. (A-F)** Elbow plots for rank selection for NMF ran on each major cell type. K was chosen as inflection point on scatter plot where rank maximizes weighted subtree metric.
